# Receptor-mediated Drp1 oligomerization on endoplasmic reticulum

**DOI:** 10.1101/190538

**Authors:** Wei-Ke Ji, Rajarshi Chakrabarti, Xintao Fan, Lori Schoenfeld, Stefan Strack, Henry N. Higgs

## Abstract

Drpl is a dynamin GTPase important for mitochondrial and peroxisomal division. Drp1 oligomerization and mitochondrial recruitment are regulated by multiple factors, including interaction with mitochondrial receptors such as Mff, MiD49, MiD51 and Fis. In addition, both endoplasmic reticulum (ER) and actin filaments play positive roles in mitochondrial division, but mechanisms for their roles are poorly defined. Here, we find that a population of Drp1 oligomers is ER-associated in mammalian cells, and is distinct from mitochondrial or peroxisomal Drp1 populations. Sub-populations of Mff and Fis1, which are tail-anchored proteins, also localize to ER. Drp1 oligomers assemble on ER, from which they can transfer to mitochondria. Suppression of Mff or inhibition of actin polymerization through the formin INF2 significantly reduces all Drp1 oligomer populations (mitochondrial, peroxisomal, ER-bound) and mitochondrial division, while Mff targeting to ER has a stimulatory effect on division. Our results suggest that ER can function as a platform for Drp1 oligomerization, and that ER-associated Drp1 contributes to mitochondrial division.

**Summary:** Assembly of the dynamin GTPase Drp1 into constriction-competent oligomers is a key event in mitochondrial division. Here, Ji *et al* show that Drp1 oligomerization can occur on endoplasmic reticulum through an ER-bound population of the tail-anchored protein Mff.

Abbreviations used in this paper: Drp1, dynamin-related protein 1; Fis1, mitochondrial fission 1 protein; INF2, inverted formin 2; KD, siRNA-mediated knock down; KI, CRISPR-mediated knock in; KO, CRISPR-mediated knock out; LatA, Latrunculin A; MDV, mitochondrially-derived vesicle; Mff, mitochondrial fission factor; MiD49 and MiD51, mitochondrial dynamics protein of 49 and 51 kDa; OMM, outer mitochondrial membrane; TA, tail-anchored.

## Introduction

Mitochondrial division plays an important role in many cellular processes, facilitating appropriate mitochondrial nucleoid distribution (Lewis et al., 2016), allowing cells to respond to changing metabolic needs (Hatch et al., 2014; Labbe et al., 2014; Mishra and Chan, 2016; Pernas and Scorrano, 2016), and contributing to selective autophagy of damaged mitochondria (Youle and van der Bliek, 2012). Defects in mitochondrial division have been linked to multiple diseases (DuBoff et al., 2013; Nunnari and Suomalainen, 2012; Vafai and Mootha, 2012).

A key component of mitochondrial division is the dynamin family GTPase Drp1. Drp1 is a cytosolic protein that is recruited to the outer mitochondrial membrane (OMM), where it oligomerizes into a spiral around the OMM (Bui and Shaw, 2013). GTP hydrolysis results in Drp1 spiral constriction, providing a driving force for mitochondrial division. Subsequent recruitment of a second dynamin GTPase, dynamin 2, appears necessary for complete membrane division (Lee et al., 2016).

A number of features suggest that mitochondrial Drp1 recruitment is a multi-step and finely-tuned process in mammals. First, mitochondrial division occurs preferentially at contact sites with endoplasmic reticulum (ER), suggesting that ER contributes components and/or signaling information to the process (Friedman et al., 2011). Second, Drp1 recruitment to mitochondria is not an all-or-none phenomenon, but rather an equilibrium process in which Drp1 oligomers dynamically assemble on mitochondria independently of signals for mitochondrial division (Ji et al., 2015). A variety of division signals may push Drp1’s on-going equilibrium toward productive oligomerization on mitochondria, including ER-mitochondrial contact, activated receptors on the OMM, cardiolipin enrichment on the OMM (Macdonald et al., 2014; Bustillo-Zabalbeitia et al., 2014), and modification of Drp1 itself (Chang and Blackstone, 2007; Chang and Blackstone, 2010; Cribbs and Strack, 2007; Friedman et al., 2011; Toyama et al., 2016). Another division signal is actin polymerization mediated by the ER-bound formin protein INF2, which stimulates division by shifting the Drp1 oligomerization equilibrium toward productive oligomerization on mitochondria (Ji et al., 2015; Korobova et al., 2014; Korobova et al., 2013). Actin’s stimulatory effect may be through direct interaction with Drp1(Hatch et al., 2016; Ji et al., 2015). Third, there are multiple Drp1 receptors on the OMM in mammals, suggesting two possibilities: 1) there are parallel pathways for Drp1 recruitment, each mediated by one of these receptors; or 2) these receptors act in a common pathway.

Protein receptors for Drp1 are necessary because, unlike other dynamin family members, Drp1 does not contain a specific lipid-binding domain. Four single-pass OMM proteins have been identified as Drp1 receptors in mammals: Mff, Fis1, MiD49 and MiD51 (Richter et al., 2015). Mff and Fis1 are tail-anchored proteins that are also found on peroxisomes, another organelle that undergoes Drp1-dependent division (Koch and Brocard, 2012; Schrader et al., 2016). In contrast, MiD49 and MiD51 contain N-terminal transmembrane domains and appear to be restricted to mitochondria (Palmer et al., 2013). Our database searches suggest that MiD49 and MiD51 are only present in vertebrates, whereas Mff is found in higher metazoans (coelomates, including arthropods and mollusks but not *C. elegans*) and Fis1 is expressed in all eukaryotes examined. Mff has consistently been found to be a key Drp1 receptor in mammals, while MiD49 and MiD51 are important in specific situations (Loson et al., 2013; Osellame et al., 2016; Otera et al., 2016; Shen et al., 2014). Though Fis1 is the sole known Drp1 receptor in budding yeast, its role in mammals is unclear (Loson et al., 2013; Osellame et al., 2016; Otera et al., 2016; Richter et al., 2015; Shen et al., 2014).

In this study, we examine Drp1 distribution among organelles in mammalian cells. Surprisingly, we find that Drp1 oligomers exist on ER, independent of mitochondrial or peroxisomal association. Populations of both Mff and Fis1 also exist on ER as punctate accumulations. Mff suppression or actin polymerization inhibition eliminates all detectable Drp1 oligomers, including the ER-bound population. We observe Drp1 accumulation at ER-bound Mff punctae, suggesting oligomeric assembly at these sites. Drp1 oligomers can transfer from ER to mitochondria or peroxisomes. Our results suggest a pathway for Drp1 oligomerization on mitochondria involving initial assembly on ER, which is dependent upon both Mff and actin.

## Results

### A sub-population of oligomeric Drp1 is bound to ER

Previously, we used an U2OS cell line stably expressing GFP-Drp1 to show that the majority (~70%) of large Drp1 “punctae” associate with mitochondria (Ji et al., 2015). These punctae likely represent Drp1 oligomers, which are clearly visible after removing the background GFP signal. To examine Drp1 localization and dynamics in more detail, we developed a GFP-Drp1 CRISPR knock-in U2OS line (called GFP-Drp1-KI), in which ~50% of the endogenous Drp1 is GFP-tagged and overall Drp1 level is similar to control cells (Fig. S1 A, B). This cell line displays similar cell growth kinetics to WT cells (Fig. S1 C), and a similar percentage of mitochondrially-associated Drp1 punctae (63%) as the stably transfected GFP-Drp1 cell line (Fig. 1 A, B).

**Figure 1.**
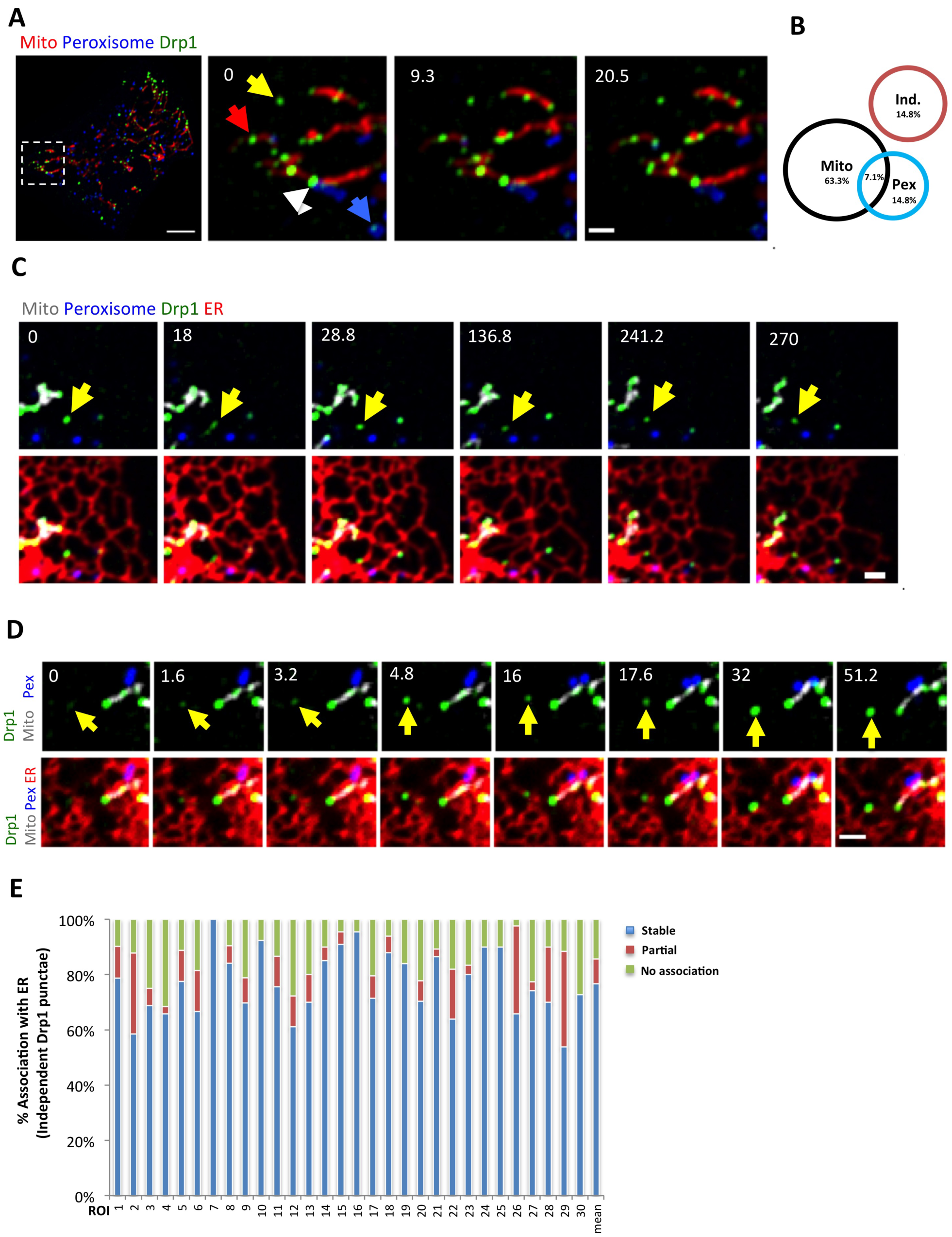
A population of Drp1 associates with ER independently of mitochondria or peroxisomes. (A) Drp1 distribution in GFP-Drp1 knock-in U2OS cells (GFP-Drp1-KI cells). *Left:* merged image of a live Drp1 KI cell transiently expressing mCherry-mito3 (red) and eBFP2-peroxisome (blue). Drp1 in green. *Right:* insets from boxed region at three time points. Yellow arrow denotes independent Drp1 puncta; blue arrow indicates peroxisome-associated Drp1; red arrow denotes mitochondrially-associated Drp1. white arrowhead denotes example of Drp1 puncta localizing at the interface of mitochondrion and peroxisome. (B) Venn diagram of Drp1 distribution in GFP-Drp1-KI cells expressing mitochondrial and peroxismal markers. Black circles denote mitochondrially associated Drp1 punctae; blue circles denote peroxisomal associated Drp1 punctae (Pex); red circles denote independent Drp1 punctae (Ind.). The percentage of Drp1 punctae in each category is average from 10 consecutive frames with 12 sec time intervals from whole-cell videos. Five cells measured (10,761 punctae). (C) Four-color imaging of a live Drp1 KI cell expressing mPlum-mito3 (Mito, gray); eBFP2-PMP20 (Peroxisome, blue); and ER-tagRFP (ER, red);Drp1 in green. Yellow arrows denote independent Drp1 puncta stably associating with ER. Video 1. (D) Time lapse montage showing *de novo* assembly of an independent Drp1 punctum (yellow arrow) on an ER tubule. Imaging as in panel C. (E) Graph depicting the degree of association between independent Drp1 punctae and ER during 2.5-min videos imaged every 1.6 sec. 30 ROIs from 25 GFP-Drp1-KI cells analyzed (1003 punctae). Mean values from ROIs: 76.7%±11.7% stable association between Drp1 punctae and ER (no apparent dissociation from ER in any frame); 8.9%±9.5% partial association; 14.4%±8.0% no association. Scale bar, 10 μm in whole cell image in (A); 2 μm in inset in (A), and in (C)&(D). Time in sec.

We examined the non-mitochondrially associated Drp1 punctae in more detail, postulating that they would be peroxisome-bound. Surprisingly, while some of these punctae are peroxisome-associated, an equal percentage (14.8%) is not associated with either mitochondria or peroxisomes, which we defined as “independent” Drp1 punctae (Fig. 1 A, B). The remaining punctae (7%) localize to areas of close association between mitochondria and peroxisomes.

We postulated that the independent population might be bound to ER. Indeed, 4-color live-cell imaging shows that a population of Drp1 punctae appears associated with ER, distinct from mitochondrial or peroxisomal populations (Fig. 1 C, Video 1). Independent Drp1 puncta can arise *de novo* from ER, maturing within 30 sec (Fig 1 D).

We quantified ER association of independent Drp1 punctae from time-lapse confocal videos, assessing stably associating punctae as those that do not separate from ER during the 2.5 min imaging time (1.6 sec frame rate). While ER occupies a significant portion of the imaging area in these cells (40.9 ± 5.9%, 22 ROIs, 2063 individual frames analyzed), there is a significantly higher percentage of independent Drp1 puncta in continual association with ER than would be expected by chance (76.7%±11.7%, Fig. 1E). Other independent Drp1 punctae are associated with ER for a portion of the imaging period (8.9%±9.5%), with most only separating for one frame. A third population of independent Drp1 punctae displays no apparent association with ER (14.4%±8.0%). Cos7 cells transiently transfected with GFP-Drp1 also display ER-associated Drp1 punctae, independent of either mitochondria or peroxisomes (Fig. S1D, E). In contrast, independent Drp1 puntae do not display appreciable association with endosomes, as judged by transferrin, Rab4b, and Rab7a markers (Fig. S2).

One possible explanation for independent Drp1 punctae is that they are actually bound to mitochondrially-derived vesicles that bud from the OMM (Soubannier et al., 2012). We tested this possibility by imaging GFP-Drp1 and the OMM protein Tom20 in live cells. No overlapping Tom20-only signal is detectable at any time point in videos (4-min, 2 sec intervals) for 15 out of 16 independent Drp1 punctae analyzed (Fig. S3). These results suggest that the majority of independent Drp1 punctae are not bound to mitochondrially-derived vesicles.

Another explanation for the existence of independent Drp1 punctae could be that they represent unfolded protein aggregates. Indeed, studies in yeast and mammals show that protein aggregates can accumulate on ER, followed by transfer to mitochondria for degradation in the mitochondrial matrix (Ruan et al., 2017; Zhou et al., 2014). While GFP-Drp1 is not over-expressed in our CRISPR-engineered cell line (Fig. S1 A-C), the GFP tag or other features of this fusion protein could result in unfolding/aggregation. To test this possibility, we examined the distribution of endogenous Drp1 punctae in relation to mitochondria, peroxisomes and ER by immunofluorescence microscopy. Similar to GFP-Drp1, a sub-set of endogenous Drp1 punctae is independent of mitochondria or peroxisomes, and 85.5%±9.7% of these independent punctae display apparent ER association (Fig. S4A, B). To confirm specificity of Drp1 immunofluorescence, siRNA suppression significantly reduces staining of all Drp1 populations (Fig. S4 A).

We examined further the effect of the GFP tag by over-expressing oligomerization-deficient mutants of Drp1 that remain monomeric or dimeric at all concentrations tested biochemically (Frohlich et al., 2013; Hatch et al., 2016). Despite being expressed at significantly higher levels than WT Drp1 in our GFP-Drp1-KI cells, these Drp1 mutants display no apparent punctae (Fig. S4 C). If the GFP tag or over-expression were causing GFP-Drp1 unfolding and aggregation, the mutants might be expected to display similar properties. We conclude that a mechanism exists for Drp1 oligomer assembly on ER.

### Transfer of Drp1 from ER to mitochondria

A range of dynamics exists for independent Drp1 punctae, with some puncta displaying little motility over a 5-min period (Fig. 1C, Video 1) while others display periods of rapid directional movement (Fig. 2A, Video 2). Independent Drp1 punctae can transfer to mitochondria (Fig. 2A, Video 2), and are ER-associated before transfer (Figure 2B, Video 3). We previously reported that most Drp1 oligomerization on mitochondria is non-productive for mitochondrial division, with only 3% of mitochondrially-associated Drp1 punctae resulting in division within a time scale of 10-min (Ji 2015). Similarly, while independent Drp1 punctae can transfer to mitochondria, division rarely occurs after these events. To increase division rate, we treated cells with ionomycin in the presence of serum, which causes a transient 4-fold increase in mitochondrial division as well as an increase in Drp1 oligomerization ((Ji et al., 2015), Fig. 3C. Fig. S5B). Upon ionomycin treatment, independent Drp1 puncta transfer to mitochondria followed by division (Fig. 2C, Video 4), with the puncta maintaining apparent association with ER during the transfer process (Fig. 2D, Video 5).

**Figure 2.**
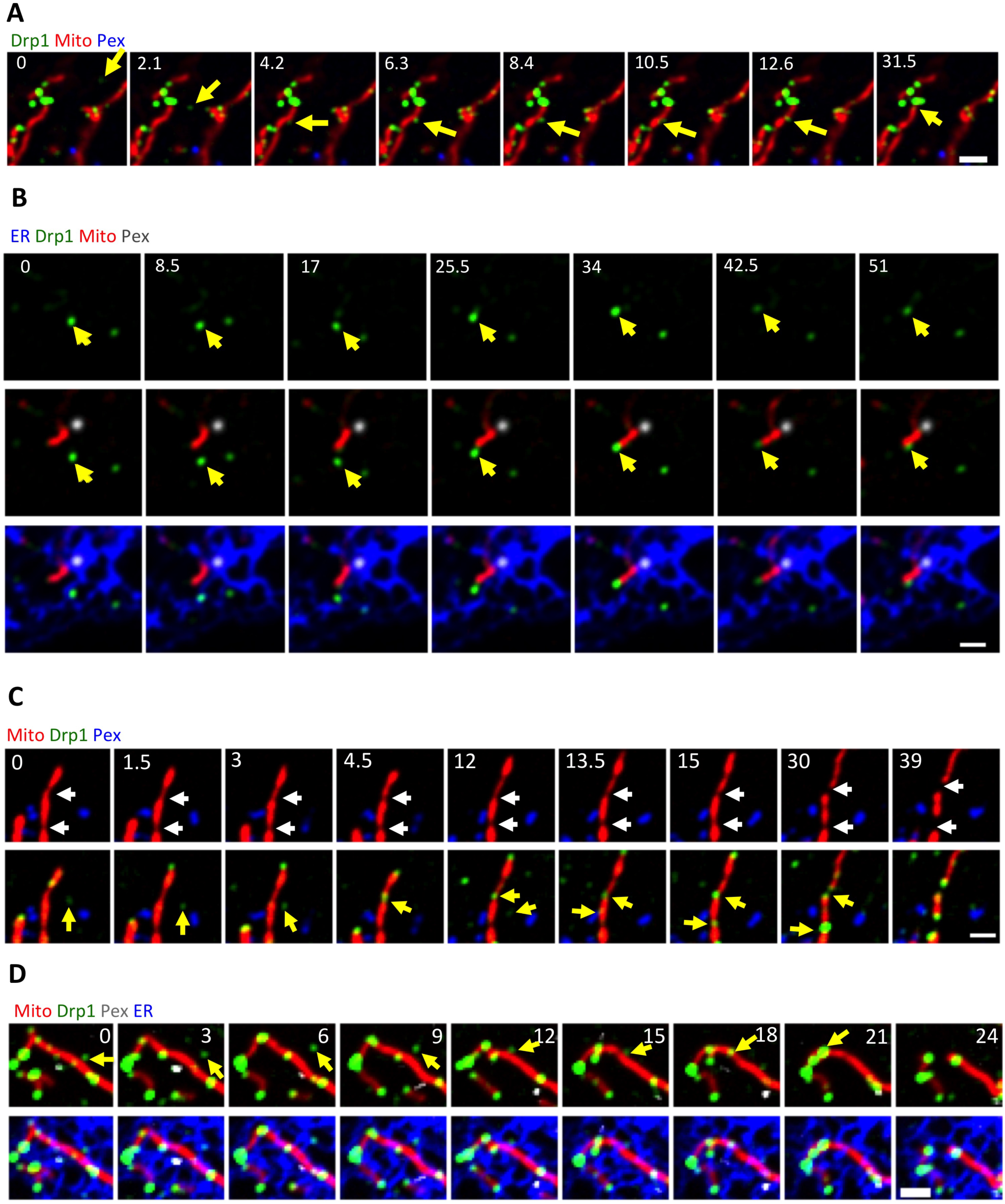
Transfer of Drp1 punctae from ER to mitochondria. (A) Three-color time lapse images of live GFP-Drp1-KI cell expressing mCherry-mito7 (mitochondria, red), eBFP2-PMP20 (peroxisome, blue) and Drp1 in green. An independent Drp1 puncta (yellow arrow) transfers to a mitochondrion and then translocates along the mitochondrion with no division in the observation time period. Video 2. (B) Four-color time lapse images of live GFP-Drp1-KI cell expressing mito-BFP(mitochondria, gray), mPlum-PMP20 (peroxisome, blue), ER-tagRFP (ER, red) and GFP-Drp1 in green. Yellow arrow denotes an ER-bound Drp1 puncta transferring to mitochondrion. Video 3. (C) Three-color time lapse images of live GFP-Drp1-KI cell expressing mCherry-mito7 (mitochondria, red), eBFP2-PMP20 (peroxisome, blue) and Drp1 in green. Two Drp1 punctae transfer to constriction sites, followed by division. Cells treated with ionomycin (4 μM) to stimulate mitochondrial division. Video 4. (D) Four-color time lapse images of live GFP-Drp1-KI cell expressing mito-BFP(mitochondria, red), mPlum-PMP20 (peroxisome, gray), ER-tagRFP (ER in blue) and GFP-Drp1 in green. Yellow arrow denotes an independent Drp1 puncta transferring to mitochondrion. Cells treated with ionomycin (4 μM) to stimulate mitochondrial division. Video 5. Scale bar: 2 μm in all images. Time in sec.

**Figure 3.**
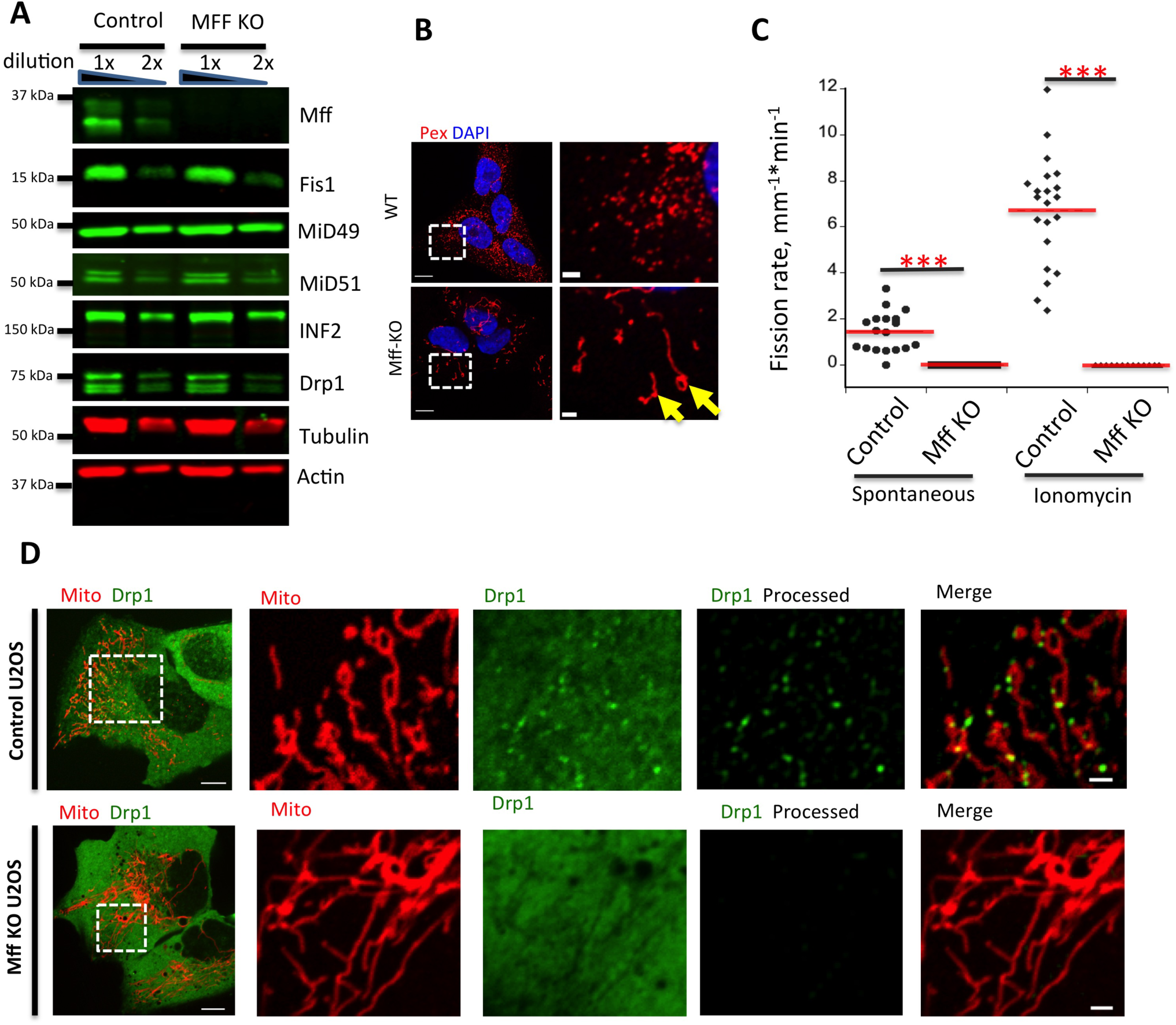
Mff knock-out U2OS cells are deficient in mitochondrial and peroxisomal division. (A) Western blotting for Mff and other mitochondrial division proteins in control and Mff KO U2OS cells. (B) Immuno-fluorescence of fixed cells stained for peroxisomes (red) and DNA (DAPI, blue). Images on right are zoomed regions. (C) Division rate quantification for both control and Mff KO U2OS cells. For the quantification of spontaneous division rate, l8 ROIs analyzed for either l2 (control) or l4 (Mff KO) cells. For quantification of ionomycin-induced division rate, 2l ROIs (control) and l3 ROIs (Mff KO) were analyzed. *** p < 0.00l by student t-test. (D) Live-cell images of control (top) or Mff KO (bottom) U2OS cells transfected with GFP-Drp1 (green) and mito-RFP (red). Right panels show ROI of selected region (boxed). Raw images shown, except for the right-most images, which are processed to reveal Drp1 punctae. Scale bars, 20 μm (left) and 2 μm (right).

### Sub-populations of Mff and Fis1 are ER-associated

We postulated that receptors on the ER membrane recruit Drp1 and enhance its oligomerization. Likely candidates for these receptors include proteins involved in mitochondrial Drp1 recruitment: Mff, MiD49, MiD51, and Fis1. There is no published evidence showing ER-bound populations of these proteins.

We first examined Mff, due to its importance for mitochondrial Drp1 recruitment in several studies (Loson et al., 2013; Osellame et al., 2016; Otera et al., 2016; Shen et al., 2014). Mff is a member of the tail-anchored family of integral membrane proteins (Gandre-Babbe and van der Bliek, 2008; Otera et al., 2010), with a C-terminal trans-membrane domain that inserts into bilayers post-translationally. We developed a CRISPR-mediated Mff knock-out (KO) cell line, that displays no detectable Mff protein but control levels of Drp1, Fis1, MiD49, MiD51, and INF2 (Fig. 3A). Similar to past studies, the Mff-KO line displays elongated peroxisomes (Fig. 3B). Mitochondrial division is almost completely eliminated in both unstimulated and ionomycin-stimulated cells (Fig. 3C). There is also a dramatic reduction in Drp1 punctae (Fig. 3D). Mff suppression by siRNA causes similar effects, including dramatic inhibition of mitochondrial division in either un-stimulated or ionomycin-stimulated cells (Fig. S5 A,B), and near-complete elimination of all Drp1 punctae in GFP-Drp1-KI cells (Fig. S4 C, Fig. S5 C). These results show that Mff is a key factor for Drp1 oligomerization in U2OS cells.

Past studies have shown Mff localization on mitochondria and peroxisomes (Friedman et al., 2011; Gandre-Babbe and van der Bliek, 2008; Otera et al., 2016; Otera et al., 2010; Palmer et al., 2013). We asked whether a sub-population of endogenous Mff was ER-bound. Using immunofluorescence microscopy in U2OS cells, endogenous Mff has a relatively uniform distribution on mitochondria and peroxisomes. In addition, there is a punctate Mff population independent of these organelles, and 89.3 ± 6.7% of these punctae associate with ER (Fig. 4A, B). This staining is specific for Mff, since Mff KD results in a dramatic reduction in all Mff populations (Fig. 4A).

**Figure 4.**
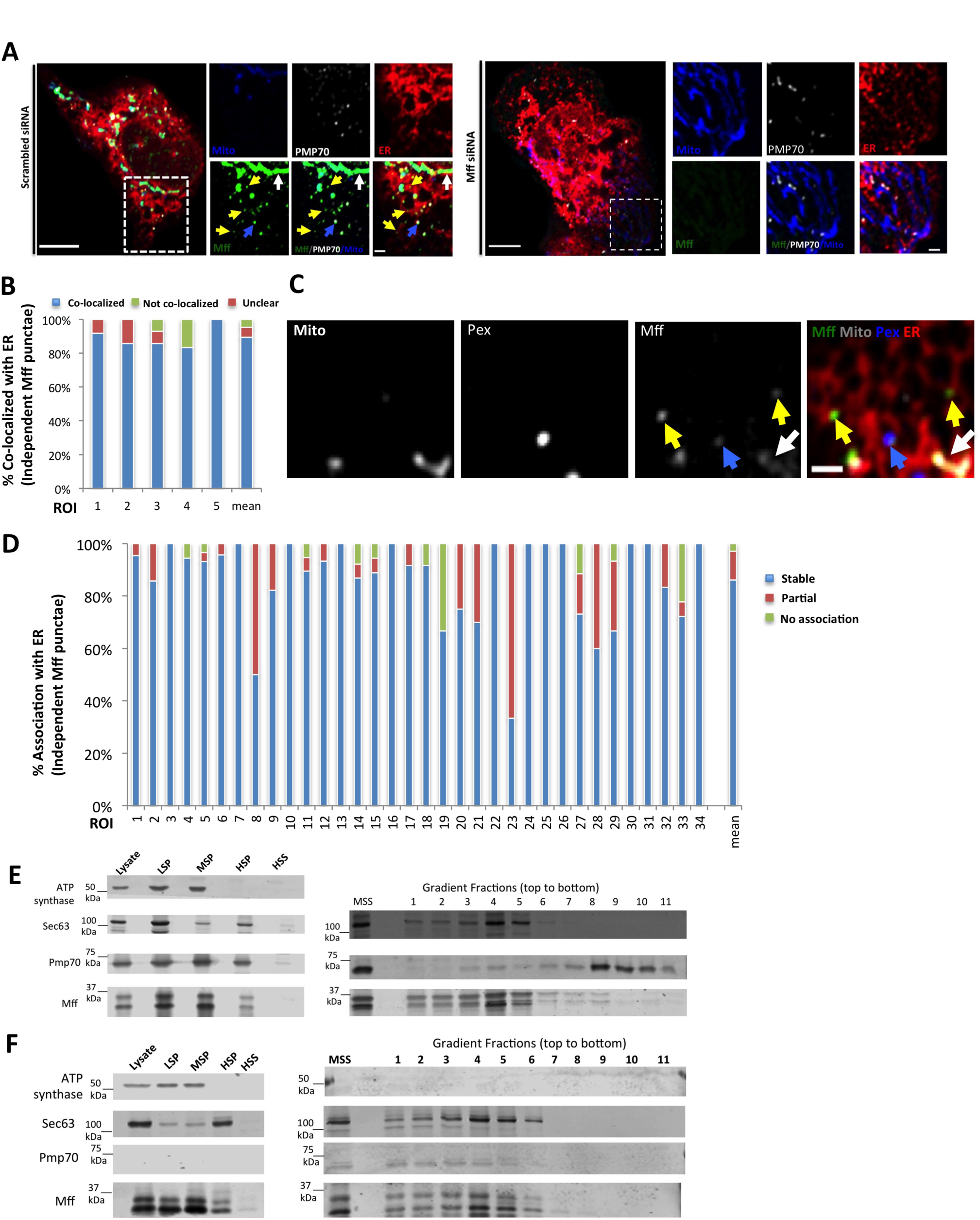
A sub-population of Mff localizes to ER. (A) Endogenous Mff localization in a fixed U2OS cell by immuno-fluorescence. Cells labeled with anti-Tom20 (mitochondria, blue), anti-PMP70 (peroxisomes, gray), anti-Mff (green); and transfected with ER-TagRFP (ER, red). Left: scrambled siRNA. Right: Mff siRNA. Independent punctae, yellow arrows. (B) Graph depicting the percentage of co-localization between independent Mff punctae and ER in U2OS cells (endogenous Mff). 54 independent Mff punctae were counted from 5 ROIs from 4 cells. Mean values from ROIs: 89.3 ± 6.7% co-localized Mff with ER, 6.0 ± 6.l% not co-localized, 4.8 ± 7.3% unclear localization. (C) Live-cell time lapse of GFP-Mff-S (green) in U2OS cell also expressing mCherry-mito3 (gray); eBFP2-peroxisome (blue); and E2-Crimson-ER (red). Yellow arrows denote independent Mff punctae associating with ER; blue and gray arrows indicate peroximal and mitochondrial Mff, respectively. Video 6. (D) Graph depicting the degree of association between independent GFP-Mff-S punctae and ER from live-cell videos as in C (2.5 min videos imaged every l.5 sec). 34 ROIs from 30 U2OS cells analyzed (44l independent Mff punctae). Mean values from ROIs: 86.1 ± l7.l% stably associated Mff punctae with ER, ll.0 ± l6.8% partially associated, 4.6 ± 9.4% not associated. (E) U2OS fractionation. Left: LSP, MSP and HSP are low, medium and high-speed pellets. HSS is highspeed supernatant. Marker proteins are: ATP synthase, mitochondria; Sec63, ER; and Pmp70, peroxisomes. Right: sucrose gradient fractionation of the MSS (medium-speed supernatant). (F) Human PEX3-deficient fibroblast fractionation, similar to U2OS fractionation. Scale bar, l0 μm in whole cell image in (B); 2 μm in inset in (B) and in (D). Time in sec.

We also examined the localization of exogenously expressed GFP-Mff in live cells. As with endogenous staining, GFP-Mff at low expression levels localizes to both mitochondria and peroxisomes. In addition, a population of independent Mff punctae is present, and 86.1 ± 17.1% of these punctae maintain continuous ER-association throughout the imaging period (Fig. 4C, D, Video 6). Mff contains four splice insert sites (Gandre-Babbe and van der Bliek, 2008). We use the variant lacking all inserts (termed Mff-S) for most investigations, but find the variant including all inserts (Mff-L) also displays this ER-localized sub-population (Fig. S6A,B)..

As a second approach to examine Mff distribution, we performed cell fractionation studies in U2OS cells. By differential centrifugation, the mitochondrial marker is confined to the low-and medium-speed pellets, whereas ER and peroxisome markers are also present in the high-speed pellet fraction (Fig. 4E). Similar to past studies (Otera 2016), Mff migrates as a ladder of bands and is present in all membrane fractions. Upon sucrose gradient fractionation of the medium-speed supernatant, ER and peroxisome markers largely separate, with a small fraction of peroxisome marker persisting in the ER fraction. Mff fractionates with the ER (Fig. 4E). These results suggest that a portion of Mff is ER-bound. To exclude the possibility that peroxisome contamination causes apparent Mff presence in the ER fraction, we used PEX3-deficient human fibroblasts, which lack mature peroxisomes (Sugiura et al., 2017). Similar to U2OS fractionation, PEX3-deficient cells contain an Mff population that fractionates with ER, and is devoid of mitochondrial and peroxisomal markers (Fig. 4F).

We also asked whether a sub-population of Fis1 is present on ER. Similar to Mff, Fis1 is a tail-anchored protein, previously reported on both mitochondria and peroxisomes (Kobayashi et al., 2007; Koch et al., 2005; Stojanovski et al., 2004; Yoon et al., 2003). By immunofluorescence analysis of endogenous protein, we observe three Fis1 populations: mitochondrial, peroxisomal and independent (Fig. 5A), with 79.9 ± 11.3% of the independent punctae displaying ER association (Fig. 5B). Fis1 depletion by siRNA strongly reduces all three of these Fis1 populations (Fig. 5A). Exogenously expressed GFP-Fis1 displays a similar population of punctae that are independent of the mitochondrial or peroxisomal Fis1 pools (Fig. 5C). Most of these independent Fis1 punctae are continually ER-associated throughout the imaging period (78.8% ± 26.9%, Fig. 5D).

**Figure 5.**
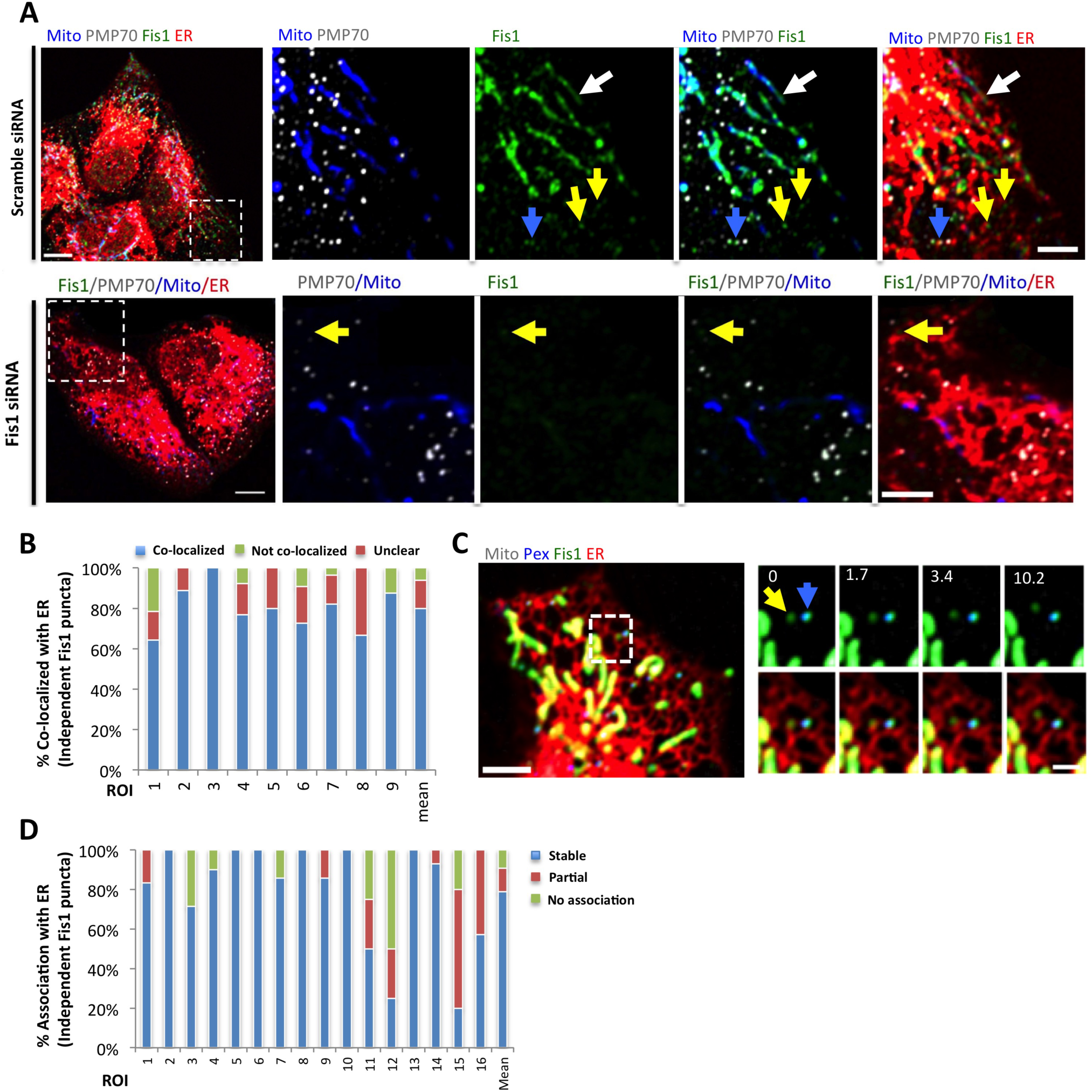
A sub-population of Fis1 localizes to ER. (A) Endogenous Fis1 localization in fixed U2OS cells by immuno-fluorescence. Cells labeled with anti-Tom20 (mitochondria, blue), anti-PMP70 (peroxisomes, gray), anti-Fis1 (green); and transfected with ER-TagRFP (ER, red). Left: scrambled siRNA. Right: Fis1 siRNA. Independent punctae, yellow arrows. (B) Graph depicting the percentage of co-localization between independent Fis1 punctae and ER in U2OS cells by immuno-fluorescence (endogenous Fis1). ll7 independent Fis1 punctae counted from 9 ROIs from 4 cells. Mean values from ROIs: 79.9 ± ll.3% co-localized Fis1 punctae with ER, 6.0 ± 7.4% not co-localized, l4.l ± l0.2% un-clear localization. (C) Live-cell time lapse of GFP-Fis1 in U2OS cell also expressing mCherry-mito3 (gray); eBFP2-peroxisome (blue); and E2-Crimson-ER (red). *Right:* individual frames from the time course of boxed region, showing independent Fis1 punctae associated with ER (yellow arrow) next to a peroxisome that is positive for Fis1 (blue arrow). (D) Graph depicting the degree of association between independent GFP-Fis1 punctae and ER from livecell videos as in C (2.5 min videos imaged every l.7 sec). l6 ROIs from l5 U2OS cells (l00 independent Fis1 punctae) analyzed. Mean values from ROIs: 78.8% ± 26.9% stably associated Fis1 punctae with ER, ll.9 ±l8.2% partially associated, 9.2 ± l4.8% not associated. Scale bar: 10 μm in whole cell image in (A) and (C); 5 μm in inset in (A); 2 μm inset in (C). Time in sec.

In contrast to Mff and Fis1, MiD49 and MiD51 contain N-terminal transmembrane domains. We examined the localization of MiD51-GFP expressed at low levels. Similar to past studies (Otera et al., 2016), MiD51 is in punctate accumulations on mitochondria, with no evidence for a peroxisomal population. There is also no evidence for a population of independent MiD51 (Fig. S6C). We conclude that both Mff and Fis1 display populations that associate with ER independently of mitochondria or peroxisomes, while MiD51 is confined to mitochondria.

### Dynamic interactions between Drp1 and Mff on ER

GFP-Mff punctae are dynamic on the ER, frequently moving and fluctuating in intensity (Fig. 4C, Video 6). We examined Mff punctae morphology and dynamics in more detail using Airyscan microscopy. As observed in the confocal images, Mff is generally distributed evenly on the surface of mitochondria and peroxisomes at low expression level, but has some regions of enrichment on both organelles (Fig. 6A, Video 7). This enrichment is particularly noticeable on peroxisomes, with one or two highly concentrated regions (Fig. 6B). The size of ER-bound Mff punctae (220 ± 56 nm, n = 19) is close to the resolution limit of Airyscan and smaller than the enriched Mff regions on peroxisomes (Fig. 6 C). Interestingly, ER-bound Mff punctae periodically appear to transfer to mitochondria (Fig. 6 A, Video 7).

**Figure 6.**
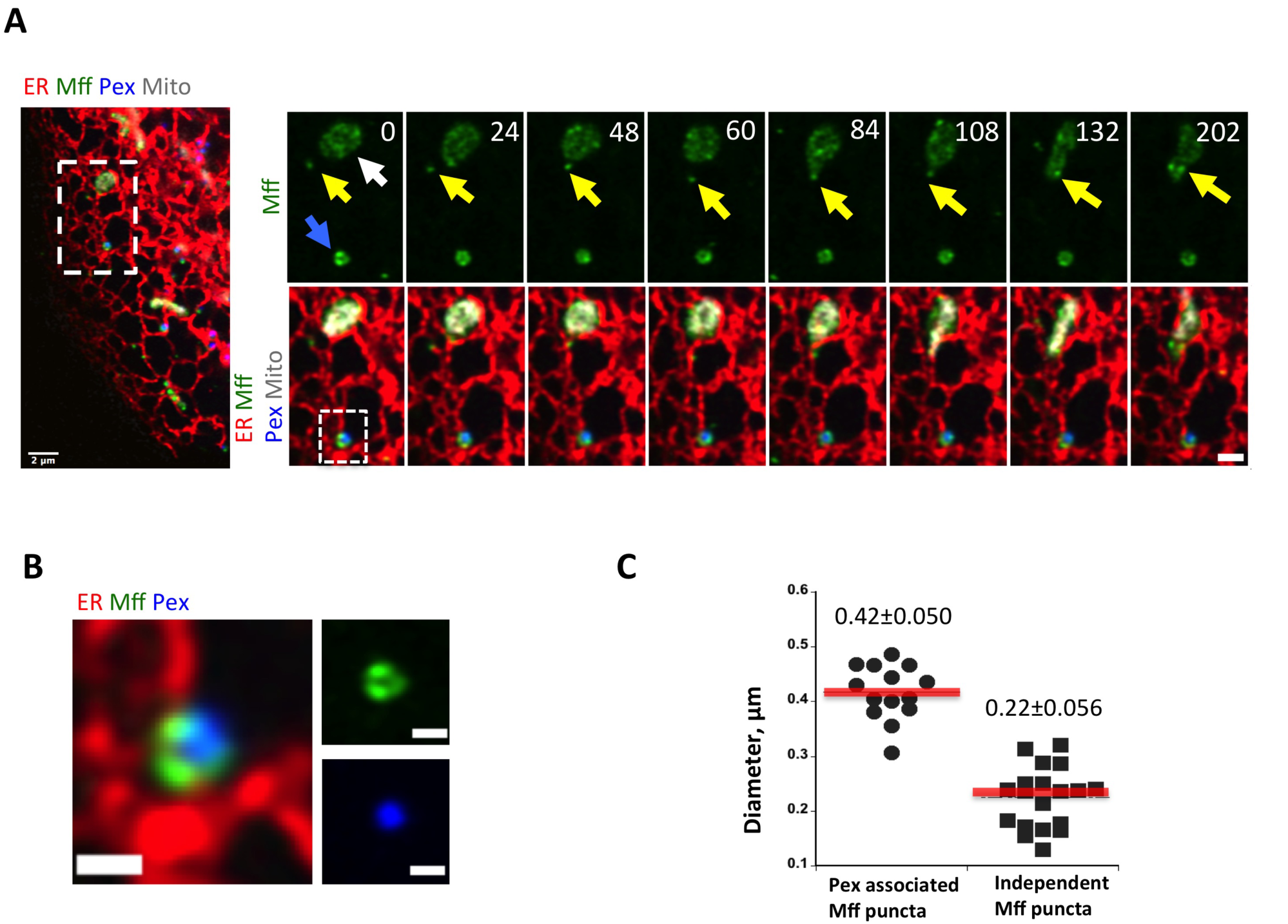
Dynamics of Mff on ER. (A) Independent Mff punctae dynamics (Airyscan microscopy time lapse). Left panel showing merged image of a live U2OS cell expressing ER-tagRFP (ER, red); GFP-Mff-S (green); eBFP2-Peroxisome (blue); and mPlum-mito3 (gray). Right panel shows a time lapse series of the inset, with independent Mff punctum associating with ER then transferring to mitochondrion (yellow arrow). Blue arrow indicates peroxisomally associated Mff and white arrow denotes mitochondrial Mff. Scale bar, 2 μm in left panel of, 1 μm in inset. Time in sec. Video 7. (B) Zoom of panel A, showing heterogeneous nature of peroxisomally-associated Mff. Scale bars, 0.5 μm in all images. (C) Dot plot showing diameter of peroxisomal Mff and independent Mff punctae from Airyscan images. 14 peroxisomal Mff (0.42±0.050 μm) and 19 independent Mff punctae (0.22±0.056) analyzed.

We next examined the relationship between Mff and Drp1 punctae on ER, using our GFP-Drp1-KI cell line transiently expressing mStrawberry-Mff at low levels. Being limited to 4-color imaging, we labeled both mitochondria and peroxisomes with BFP, and labeled ER with an E2-crimson marker (Fig. 7A, Video 8). From quantification of live-cell time-lapse images, ~70% of the ER-bound Drp1 and Mff punctae co-associate for the entirety of the 3-min imaging period (98 of 140 Mff punctae associated with Drp1, 84 of 140 Drp1 punctae associated with Mff). There are also instances of Drp1 appearance from previously existing Mff punctae (Fig. 7A, Video 8), suggesting that ER-bound Mff punctae are sites of Drp1 oligomerization. Interestingly, the number of independent Mff punctae decreases ~ 4-fold upon Drp1 suppression by siRNA, either when analyzing GFP-Mff in live cells (4.5-fold decrease, Fig. 7B, C), or endogenous Mff by immunofluorescence (3.9-fold decrease, Fig. 7D).

**Figure 7.**
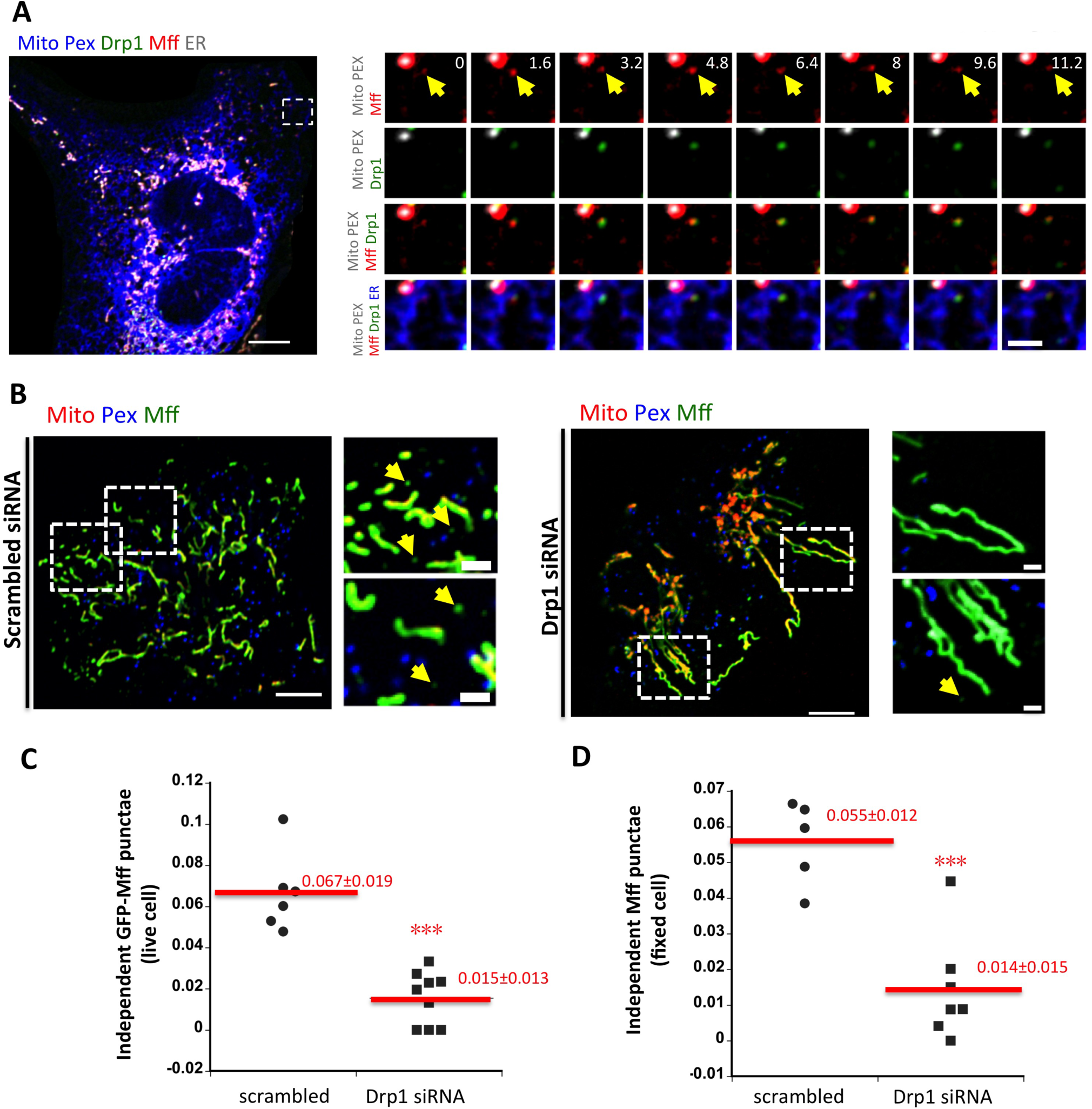
Association between Drp1 and Mff on ER. (A) Left: Merged confocal image of a live GFP-Drp1-KI cell expressing mito-BFP(gray), eBFP2-peroxisome (gray), mStrawberry-Mff-S(red) and pLVX-E2-Crimson-ER (blue). Drp1 in green. *Right:* Time lapse confocal images of boxed region show example of a Drp1 puncta maturing from an independent Mff puncta (yellow arrows). Video 8. (B) Independent Mff punctae in scramble siRNA treated (left) and Drp1 siRNA treated cells (right). *Left:* merged image of live U2OS cells transiently expressing GFP-Mff-S(Mff, green), eBFP2-PMP20 (Pex, blue) and mCherry-mito7 (Mito, red). *Right:* insets from boxed regions in whole cell image. Yellow arrows denote independent Mff punctae. (C) Density of independent Mff punctae in control and Drp1 siRNA-treated U2OS cells, quantified from live cell images of GFP-Mff as (B). Units, number of independent Mff punctae per μm^2^ in the ROI. 368 independent punctae from nine control cell ROIs and 106 punctae from nine Drp1 KD cell ROIs. *** denotes p value < 0.0001 by student t-test. (D) Density of independent Mff punctae in control and Drp1 siRNA-treated U2OS cells, quantified from fixed cell immuno-fluorescence of endogenous Mff. Units, number of Mff punctae per μm^2^ in ROI. 643 independent punctae from five control cell ROIs and 153 puncta from seven Drp1 KD cell ROIs. *** denotes p value < 0.0005 by student t-test. Scale bar: 10 μm in whole cell images; 2 μm in insets. Time in sec

### ER-localized Mff enhances mitochondrial division rate

To test the functional significance of ER-targeted Mff, we designed a rapamycin-inducible system in which Mff lacking its transmembrane domain could be targeted to either mitochondria or ER, using the targeting sequences of AKAP1 and Sac1, respectively (Fig. 8A, (Csordas et al., 2010)). A similar approach has been used to target Mff to lysosomes (Liu and Chan, 2015). Rapamycin treatment results in rapid Mff translocation from cytosol to mitochondria in Mff-KO U2OS cells (Fig. 8B). Rapamycin-induced translocation to ER is also rapid, but with some Mff still present in cytoplasm (Fig. 8C). We used this system to test the effect of targeting Mff to specific locations (mitochondria alone, ER alone, or to both mitochondria and ER) on mitochondrial division rate in Mff-KO cells. While either mitochondrial or ER targeting causes partial rescue, targeting Mff to both organelles brings the mitochondrial division rate back to the level of control cells (Fig. 8D). The enhanced effect of expressing both mitochondrial and ER targeting signals is not due to increased expression of Mff or of Drp1 (Fig. 8E). These results suggest that ER targeting of Mff has a stimulatory effect on mitochondrial division.

**Figure 8.**
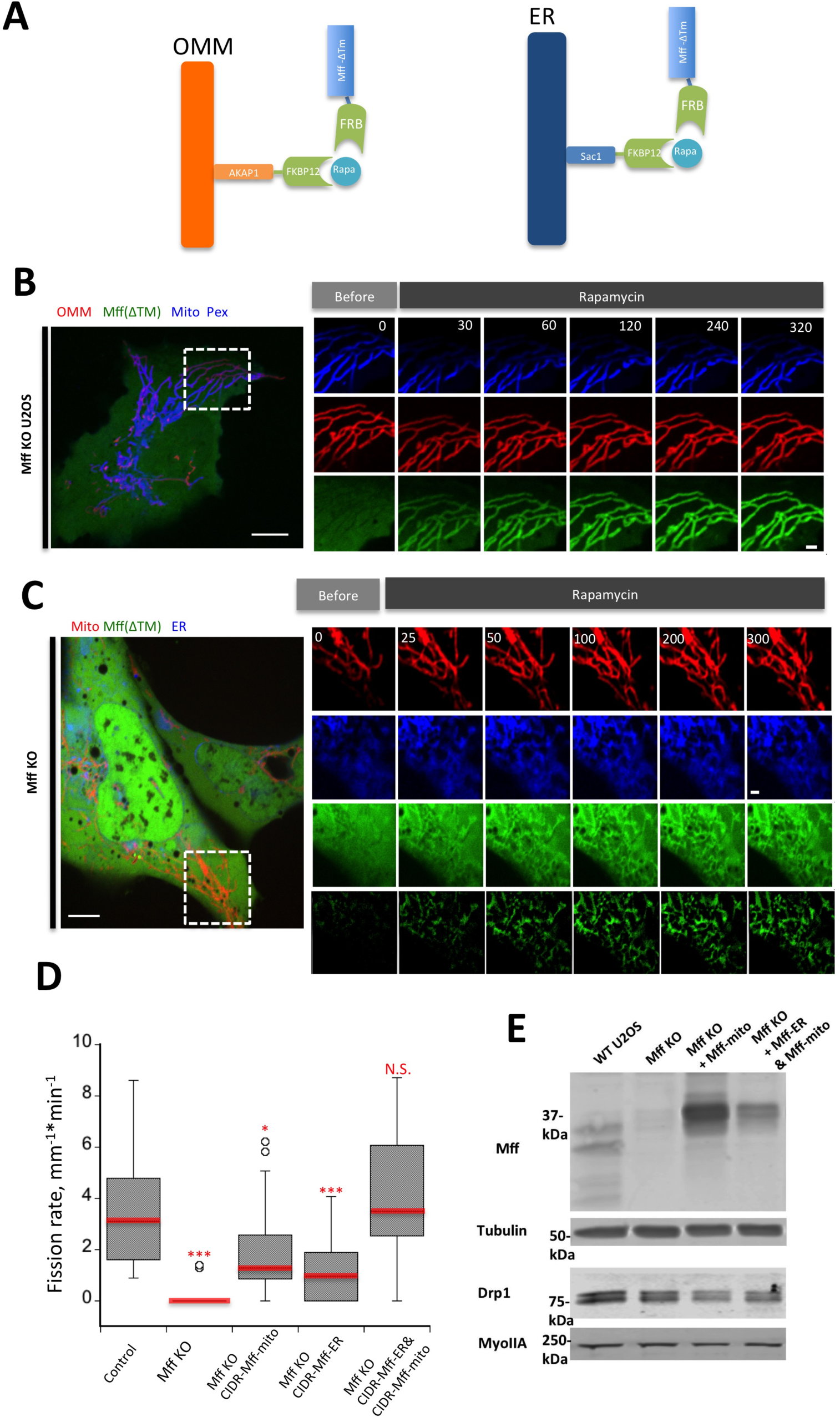
ER-targeted Mff facilitates mitochondrial division. (A) Schematic cartoon of rapamycin-induced Mff recruitment either to OMM (left) or ER (right). “Mff” refers to the cytoplasmic portion of Mff-S. (B) Dynamics of GFP-Mff-FRB translocation to mitochondria upon rapamycin treatment in Mff KO cells. Live cell images of cell transfected with AKAP-FKBP12 (red), GFP-Mff-Cyto-FRB (green), eBFP2-PMP20 (peroxisomes, blue) and mitoBFP (Mitochondria, blue). Rapamycin (final concentration: 10μM) added at time 0. (C) Dynamics of GFP-Mff-Cyto translocation to ER upon rapamycin in rapamycin treatment in Mff KO cells. Live cell images of cells transfected with Sac1-FKBP12 (ER, blue), GFP-Mff-Cyto-FRB (green), and mCherry-mito7 (mitochondria, red). The lower green panel represents GFP-MFF-CytoFRB signal that has been thresholded to remove the cytoplasmic signal. Rapamycin (final concentration: 10μM) added at time 0. (D) Rapamycin-induced mitochondrial division rates in control U2OS cells (l6 ROIs from l5 cells); Mff KO cells (2l ROIs from 2l cells) (P=0.0000l, ***); Mff KO cells transfected with mitochondria-targeted Mff (34 ROIs from 30 cells) (P=0.0l79,*); Mff KO cells transfected with ER-targeted Mff (20 ROIs from l7 cells) (P=0.0049,***); or Mff KO cells transfected with both mitochondria‐ and ER-targeted Mff (34 ROIs from 30 cells)(P=0.4l8l). Statistical analysis based on comparison to control cells by student t-test. (E) Western blot showing Mff and Drp1 expression levels in WT cells, Mff-KO cells, and Mff-KO cells transfected with either the Mff-FRB construct + the mitochondrially-targeted FKBPl2 construct (Mff KO + Mff-mito) or the Mff-FRB construct + the mitochondrially-targeted FKBPl2 construct + the ER-targeted FKBPl2 construct (Mff KO + Mff-ER & Mff-mito). Tubulin and myosin IIA are loading controls. Endogenous Mff runs as a doublet below 37 kDa, whereas the Mff-FRB construct runs at the 37 kDa marker. Scale bar: l0 μm in whole cell images in (B, C);2 μm in insets in (B, C). Time in sec.

### ER-associated Drp1 oligomers are dependent on INF2-mediated actin polymerization

In a previous study (Ji et al., 2015), we found that ionomycin enhances Drp1 maturation on mitochondria. To test whether ionomycin can trigger ER-associated Drp1 maturation as well, we tracked independent Drp1 punctae upon ionomycin treatment. Ionomycin significantly increases both the number (Fig. 9, Video 9) and size (Fig. 10A, B) of independent Drp1 punctae.

**Figure 9.**
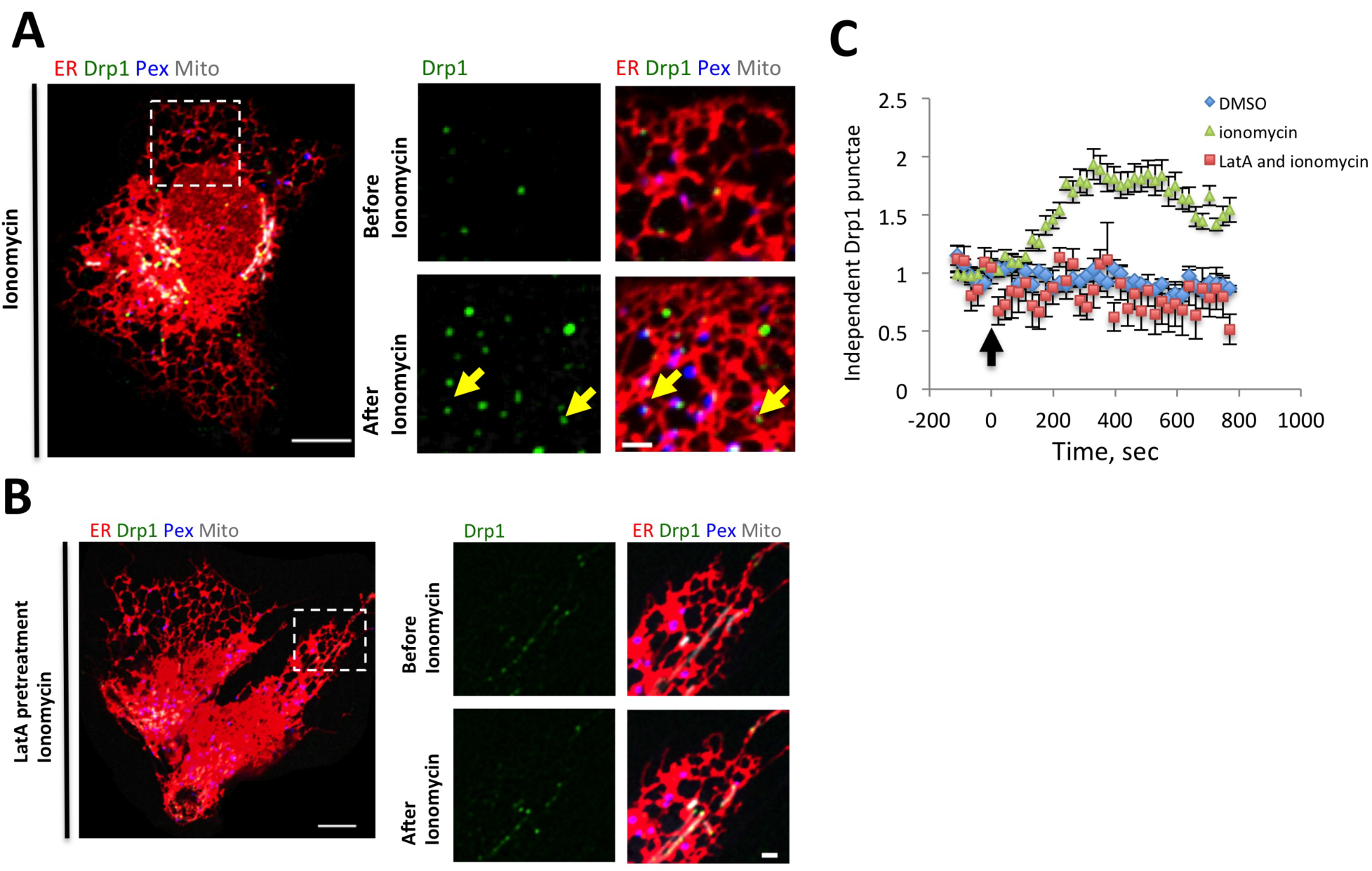
Actin-dependent oligomerization of ER-associated Drp1 punctae. (A) Left: merged image of a live GFP-Drp1-KI cell before ionomycin treatment, transiently expressing mPlum-mito3 (gray), eBFP2-peroxisome (blue), and ER-tagRFP (ER, red). Drp1 in green. Right: inset from boxed region before (top) and after (bottom) ionomycin treatment (4 μM, l0 min). Yellow arrows denote independent Drp1 maturing upon ionomycin treatment. Video 9. (B) Similar experiment as in A, except cells were pre-treated for l0min with l μM LatA. Video l0. (C) Quantification of independent Drp1 punctae number in response to vehicle treatment (DMSO), ionomycin treatment and LatA pre-treatment followed by ionomycin treatment. 6 ROIs from 6 DMSO treated cells, l6 ROIs from l4 ionomycin treated cells, and 8 ROIs from 6 LatA pre-treated/ionomycin treated cells. Punctae per ROI normalized to l at time of ionomycin addition. Error bar, S.E.M. Arrow indicates time point where ionomycin was added during imaging (time 0). Scale bar: l0 μm in left panels; 2 μm in right panels. Time in sec.

**Figure 10.**
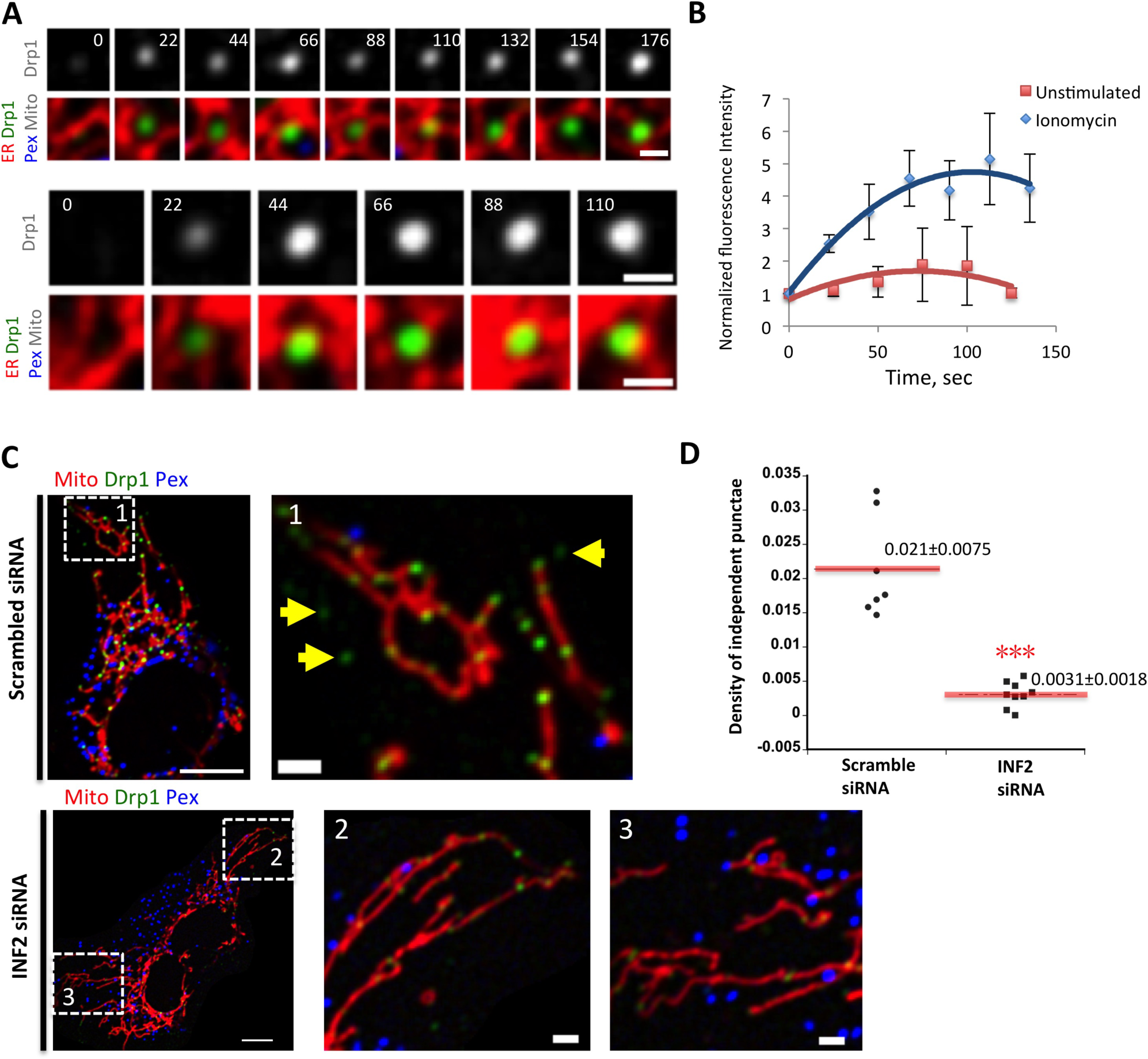
Maturation of existing independent Drp1 punctae upon ionomycin stimulation. (A) Two examples of independent Drp1 punctae maturation in response to ionomycin. Time-lapse images of live GFP-Drp1-KI cell as in Fig. 9A. Time indicates sec after ionomycin treatment. Fluorescence intensity levels modulated uniformly across timecourse so that final fluorescence is in linear range (resulting in time 0 fluorescence being undetectable as displayed). Scale bars, l μm. Time in sec. (B) Quantification of mean independent Drp1 punctum intensity in un-stimulated or ionomycin treated conditions. Seven independent Drp1 punctae from un-stimulated cells and eight independent Drp1 punctae from ionomycin treated analyzed. Error bars, S.D. (C) Effect of INF2 KD on independent Drp1 punctae in GFP-Drp1-KI cells transfected with mCherry-mito7 (mitochondria, red) and eBFP2-peroxisome (blue). Drp1 in green. Top is control siRNA, bottom is INF2 siRNA. *Right:* zoomed images of boxed regions indicated by numbers. Scale bar: l0 μm in left panels; 2 μm in right panels (insets). (D) Quantification of independent Drp1 punctae density in control (scrambled siRNA) and INF2 siRNA cells. l74 independent punctae from seven control cells, 45 independent punctae from nine INF2 siRNA cells. Density expressed as number of independent Drp1 punctae per area of ROI (in μm^2^). *** denotes p < 0.00l by student’s t-test.

Our previous studies also showed that mitochondrially-bound Drp1 oligomers are significantly decreased by actin polymerization inhibitors (Korobova et al., 2013) and that actin polymerization inhibitors block the ionomycin-induced increase in Drp1 oligomerization (Ji et al., 2015). We tested the effect of Latrunculin A (LatA), an actin polymerization inhibitor, on ER-bound Drp1 oligomers. Pre-treatment for 10 min with LatA causes a significant reduction in all Drp1 punctae prior to ionomycin treatment, and a near-complete block of independent Drp1 punctae maturation upon ionomycin treatment (Fig. 9, Video 10).

We have shown that the formin INF2 is required for actin polymerization leading to efficient mitochondrial division (Korobova et al., 2013) as well as mitochondrial accumulation of oligomeric Drp1 (Ji et al., 2015). The isoform of INF2 responsible for these effects is tightly bound to ER (Chhabra et al., 2009), suggesting that it could also play a role in ER-bound Drp1 oligomerization. We therefore tested whether INF2 played a role in independent Drp1 punctae accumulation. Suppression of INF2 by siRNA causes a 6.8-fold decrease in independent Drp1 punctae (Fig. 10C, D). These results indicate that INF2-mediated actin polymerization is necessary for ER-associated Drp1 oligomerization.

## Discussion

A major finding in this work is the identification of dynamic sub-populations of Drp1, Mff and Fis1 on ER, distinct from the mitochondrial and peroxisomal populations of these proteins. An earlier study suggested that Drp1 could localize to ER (Yoon et al., 1998), but this study did not include mitochondrial or peroxisomal markers so specific localization to ER is unclear. There has been no previous identification of ER-bound sub-populations of any wild-type Drp1 receptor. We carefully examined previous publications for evidence of such localization for Mff (Friedman et al., 2011; Gandre-Babbe and van der Bliek, 2008; Otera et al., 2016; Palmer et al., 2013) or Fis1 (Kobayashi et al., 2007; Koch et al., 2005; Stojanovski et al., 2004; Yoon et al., 2003). Most of these studies do not stain for both peroxisomes and mitochondria, but in two studies using both markers we find evidence for Mff (Palmer et al., 2013) and Fis1 (Kobayashi et al., 2007) punctae that are not bound to either organelle. The low abundance of these independent punctae, and their low intensities compared to both the mitochondrial and peroxisomal pools, could explain why this population has not been identified previously. Interestingly, a recent proteomic study identified an apparent ER-linked pool of Mff by proximity ligation (Hung et al., 2017), which could be an ER-bound population but could alternately represent a population at ER-mitochondrial contact sites.

Mff and Fis1 are tail-anchored (TA) proteins that are inserted into membranes post-translationally. TA proteins are found in essentially all cellular membranes, including ER, mitochondria and peroxisomes. Insertion mechanisms for ER-based TA proteins are best understood, with the GET/TRC40 complex being an important pathway (Denic et al., 2013; Mateja et al., 2015; Schuldiner et al., 2008; Stefanovic and Hegde, 2007), and the recently identified SND pathway being an alternate route (Aviram et al., 2016). At present, the pathways controlling TA protein targeting to mitochondria or peroxisomes are less well understood, with evidence for three routes: 1) protein-free insertion (Krumpe et al., 2012), 2) protein-mediated insertion (Yagita et al., 2013), and 3) delivery from ER (Lam et al., 2010; Schuldiner et al., 2005; van der Zand et al., 2010).

The presence of Mff and Fis1 on all three membranes does not clarify their delivery mechanisms, but their wider distribution suggest mechanisms that would lead to both ER and mitochondrial insertion. Interestingly, one study (Stojanovski et al., 2004) showed that mutagenesis of two C-terminal lysines in mammalian Fis1 caused a shift in its localization from mitochondria to ER, which might suggest mitochondrial localization signals in the C-terminus similar to findings for other proteins (Horie et al., 2002). It is also interesting that Mff is undetectable in the peroxisomes present in the “light” membrane fraction of U2OS cells (Fig. 4E), suggesting these peroxisomes are different from those in the heavier membrane fractions.

While we provide evidence that the ER-localized pool of Mff acts in mitochondrial division, there are other possible explanations for Mff’s presence on ER. First, a portion of the ER pool might represent a transient intermediate in Mff’s bio-synthetic pathway, in which it is first inserted into the ER membrane then transferred to the OMM. Alternately, a portion ER-bound Mff and Fis1 might represent mis-localized protein that is subsequently sorted to the OMM by a secondary sorting mechanism. These possibilities are not mutually exclusive with the existence of a functional pool of ER-localized Mff. Better understanding of targeting mechanisms for Mff is required, including pulse-chase localization studies to determine whether an ER intermediate exists.

Recent studies in budding yeast and mammals show that aggregates of misfolded protein can bind ER, then move to the mitochondrial matrix for proteolysis (Ruan et al., 2017; Zhou et al., 2014). However, several lines of evidence strongly suggest that the ER-bound punctae of Drp1, Mff and Fis1 observed here are not protein aggregates. First, in all three cases we observe these punctae using immunofluorescence for endogenous proteins, arguing against over-expression artifact. Second, GFP-fusions of non-oligomerizable Drp1 mutants do not display ER-bound punctae, even when expressed at significantly higher levels than wild-type GFP-Drp1. Third, ER-bound Drp1 punctae are virtually absent in the following conditions: Mff knock-out, actin polymerization inhibition, and suppression of the actin polymerization factor INF2. All of these conditions inhibit mitochondrial division but are not known to be related to aggregated protein responses. LatA treatment reduces the number of independent Drp1 punctae within 10 min, demonstrating the dynamic nature of this population.

Another possibility is that independent Drp1 or Mff punctae represent mitochondrially-derived vesicles (MDVs) containing OMM but not IMM. Our imaging of Drp1 and the OMM protein Tom20 suggest that this is not the case, as we observe no consistent co-localization. Even so, MDVs can have heterogeneous composition (Soubannier et al., 2012), which leaves the possibility open that the independent punctae are bound to a specific MDV sub-type. Since the majority of independent Drp1, Mff and Fis1 punctae track tightly with ER in live-cell imaging, any MDV would likely be associated with ER in this case.

One possible functional role of ER-assembled Drp1 is in mitochondrial division. In support of this function, 1) we observe transfer of Drp1 punctae from ER to mitochondria; 2) we observe mitochondrial division following ER-to-mitochondrial Drp1 transfer; and 3) in Mff-KO cells, targeting Mff to both ER and mitochondria is more efficient in rescuing mitochondrial division than is targeting to either ER or mitochondria alone. There are uncertainties in this correlation. Limitations of confocal microscopy in both spatial and temporal resolution make it difficult to be certain of direct ER-to-mitochondrial Drp1 transfer. In addition, there is a significant amount of ER-to-mitochondrial Drp1 transfer that does not result in mitochondrial division. To observe mitochondrial division following ER-to-mitochondrial Drp1 transfer, we stimulate division frequency with the calcium ionophore ionomycin. However, mitochondrial division in general occurs at low frequency, and the vast majority of mitochondrially-bound Drp1 punctae in general are non-productive for mitochondrial division (Ji et al., 2015), suggesting that Drp1 oligomerization is in dynamic equilibrium independent of mitochondrial division.

From our findings, we propose a working model that includes a role for ER in Drp1 oligomerization and recruitment prior to interaction with mitochondria or peroxisomes. The combination of ER-bound Mff and INF2-mediated actin polymerization on ER serves as an initiation site for recruitment of Drp1 oligomers. These Drp1 oligomers can be transferred to mitochondria or peroxisomes upon ER contact, where they can serve in the assembly of mitochondrially-bound Drp1 oligomers capable of mitochondrial division. This Drp1 transfer can occur without transfer of the receptors themselves, although we have observed movement of Mff punctae between ER and mitochondria. Further study of these dynamics is needed.

Assembly on ER is likely to be only one component of Drp1’s oligomeric equilibrium, in addition to direct assembly on mitochondria or peroxisomes. Another possibility is that the independent Drp1 punctae observed here represent a minor proportion of all ER-bound Drp1 oligomers, with the vast majority being assembled on ER at ER-mitochondrial contact sites. Due to the close proximity of ER-mitochondrial contact sites (Csordas et al., 2010), imaging Drp1 transfer at these sites is challenging by current live-cell techniques.

Our work adds another layer to the understanding of roles for mammalian Drp1 receptors (Mff, Fis1, MiD49 and MiD51) in mitochondrial and peroxisomal division. The current picture is somewhat murky, with recent knock-down/knock-out studies in several cell lines providing largely overlapping but at times conflicting results (Loson et al., 2013; Osellame et al., 2016; Otera et al., 2016; Shen et al., 2014). One feature of clear agreement is that neither MiD49 nor MiD51 localizes to peroxisomes, and neither participates in peroxisomal division (Otera et al., 2016; Palmer et al., 2013). Another common theme is that MiD49 and MiD51 are at least partially redundant with each other, and have the capability of acting independently of Mff (Loson et al., 2013; Osellame et al., 2016; Otera et al., 2016; Palmer et al., 2013). Most studies find the role of Fis1 in Drp1 recruitment and mitochondrial/peroxisomal division to be minor at best, although one study finds more significant effects (Shen et al., 2014). Deletion of Mff typically has the most dramatic effects on both Drp1 recruitment and mitochondrial division, but one study finds that MiD49/51 deletion has comparable effects (Osellame et al., 2016). The differing results may be partly due to cellular context. In mitophagy, for example, Fis1 might play a role in Drp1 recruitment downstream of Mff (Shen et al., 2014). During apoptosis, MiD49 and MiD51 have roles in cristae remodeling (Otera et al., 2016), although other Drp1 receptors clearly function in apoptosis as well (Osellame et al., 2016).

We make three important findings on Drp1 receptors in this work. First, Mff is of fundamental importance in U2OS cells, since either siRNA-mediated suppression or CRISPR-mediated knock-out strongly reduce both Drp1 oligomerization and mitochondrial division. Second, U2OS cells have ER-bound populations of both Mff and Fis1. Third, the majority of ER-bound Drp1 punctae co-localize with Mff. These populations appear to be co-dependent, with reduction of either Drp1 or Mff reducing punctae of the other protein on ER. Presumably, Drp1 oligomerization recruits additional Mff from the bulk ER.

One open question concerns why there are so many potential mechanisms for regulating Drp1, including: multiple receptors (MiD49, MiD51, Fis1), Drp1 post-translational modification (Chang and Blackstone, 2007; Chang and Blackstone, 2010; Cribbs and Strack, 2007), cardiolipin enrichment on the OMM (Macdonald et al., 2014; Bustillo-Zabalbeitia et al., 2014), actin polymerization (Korobova et al., 2013; Li et al., 2014; Ji et al., 2015; Moore et al., 2016), and ER-mitochondrial contact (Friedman et al., 2011). Do these mechanisms operate in concert or independently? Given that Drp1 oligomer assembly and disassembly are constantly in flux on mitochondria (Ji et al., 2015), the answer could be “both”. A critical threshold of Drp1 oligomerization and mitochondrial recruitment is necessary, regardless of the means by which oligomerization/recruitment are activated. In this model, a variety of combinations of these activators can lead to the final outcome of division-productive Drp1 oligomerization. Other aspects of Drp1-mediated force generation may be similarly nuanced (Ramachandran, 2017). Importantly, the ER-based recruitment of Drp1 oligomers represents only one of these activation mechanisms, and its loss may be compensated by up-regulation of the other mechanisms. An additional step may be recruitment of dynamin 2 late in the process (Lee et al., 2016), which would be subject to its own regulation.

This study extends our findings on the role of actin in mitochondrial division by showing actin polymerization is necessary for initiation and growth of ER-bound Drp1 oligomers. We have proposed direct binding of Drp1 to actin filaments as a potential mechanism for increasing productive Drp1 oligomerization (Hatch et al., 2016; Ji et al., 2015). The presence of the formin INF2 on ER (Chhabra et al., 2009) and its importance for Drp1 recruitment to mitochondria (Ji et al., 2015; Korobova et al., 2013) suggest that INF2-mediated actin polymerization on ER, in conjunction with Mff on ER, might mediate ER-based Drp1 oligomerization. Actin polymerization has been implicated in mitochondrial division in many contexts (De Vos et al., 2005; Duboff et al., 2012; Moore et al., 2016) and additional actin-binding proteins on mitochondria (Manor et al., 2015) and in cytosol (Li et al., 2015; Moore et al., 2016) have been implicated. It will be interesting to elucidate if and how these proteins work together in this process.

Given the presence of Drp1 oligomers on ER, and its role in constriction of mitochondria and peroxisomes, it is tempting to speculate that Drp1 might mediate some aspect of ER membrane dynamics. Past studies have suggested that dominant-negative Drp1 mutants change ER structure (Pitts et al., 1999). While we have occasionally observed Drp1 punctae at sites of ER tubule breakage (not shown), these instances are rare. Nevertheless, the presence of Drp1, Mff and Fis1 on ER expands mechanistic possibilities for membrane dynamics in general.

## Materials and Methods

### Plasmids and siRNA oligonucleotides

mCherry-mito-7 was purchased from Addgene (#55102), and consists of the mitochondrial targeting sequence from subunit VIII of human cytochrome C oxidase N-terminal to mCherry. mito-BFP construct were previously described(Friedman et al., 2011), and consist of amino acids 1-22 of S. cerevisiae COX4 N-terminal to BFP. Tom20-mCherry was previously described in(Ji et al., 2015). eBFP-Peroxisome was constructed by replacing the CFP sequence of CFP-Peroxisome containing peroxisomal targeting signal 1 (PTS1) (Addgene #54548) with eBFP2, cut from eBFP2 ‐Mito7 (Addgene #55248) with BsrGI/BamHI. mPlum-mito3 was purchased from Addgene (#55988). ER-tagRFP was a gift from Erik Snapp (Albert Einstein College of Medicine, New York City), with prolactin signal sequence at 5' of the fluorescent protein and KDEL sequence at 3'. pEF.myc.ER-E2-Crimson was purchased from Addgene (#38770). Mff-S and MiD51 cloned by reverse transcriptase-PCR from RNA isolated from HEK293 cells and cloned into eGFP-C1 (Mff) or eGFP-N1 (MiD51) vectors (Clontech Inc). GFP-Mff-L was purchased from Addgene (# 49153). mStrawberry-Mff-S was constructed by replacing GFP with mStrawberry using SalI/BamHI. MiD51-mStrawberry was constructed by cutting MiD51 from MiD51-GFP and pasting into mStrawberry-N1 vector using Bgl II/BamH1. GFP-Fis1 was a gift from Mike Ryan (Monash University, Melbourne, Australia). mStrawberry-Rab4b and mStrawberry-Rab7a were gifts from Mitsunori Fukada (Tohoko University, Sendai, Japan) (Matsui et al., 2011). In our nomenclature for Mff isoforms, Mff-S corresponds to isoform 8 (no alternately spliced exons) and Mff-S corresponds to isoform 1 (containing all alternately spliced exons) from (Gandre-Babbe and van der Bliek, 2008). Drp1 mutants that maintain the monomeric (K642E) or dimeric (K401-404A) states were described in (Hatch et al., 2016). Rapamycin-inducible constructs include the following. Mitochondrial targeting construct: amino acids 1-3l of mouse AKAP1 fused to FKBP12. ER-targeting construct: C-terminal sequence of human/mouse Sacl fused to FKBP12. Both mitochondrial‐ and ER-targeted FKBP12 constructs were generous gifts of Gyorgy Hajnoczky (Csordas et al., 2010). GFP-Mff inducibly-targetable construct: the cytoplasmic region (amino acids 1-197) of human Mff-S fused to GFP on the N-terminus and FRB on the C-terminus.

Oligonucleotides for human Mff siRNA were synthesized by Qiagen against target sequence 5’-ACCGATTTCTGCACCGGAGTA-3’. Oligonucleotides for MiD51 were synthesized by Qiagen against sequence 5’-CAGTATGAGCGTGACAAACAT ‐3’ (siRNA#1), and 5’-CCTGGTCTTTCTCAACGGCAA ‐3’ (siRNA#2). Oligonucleotides for MiD49 were synthesized by Qiagen against sequence 5’-TTGGGCTATGGTGGCCATAAA-3’ (siRNA#1), and 5’-CTGCTGAGAGAAGGTGACTTA-3’ (siRNA#2). Oligonucleotides for Fis1 were synthesized by IDT against target sequence 5’-GUACAAUGAUGACAUCCUAAAGGC-3’ (siRNA#1), and 5’-ACAAUGAUGACAUCCGUAAAGGCAT-3’ (siRNA#2). Oligonucleotides for human total INF2 siRNA were synthesized by IDT Oligo against target sequence 5’-GGAUCAACCUGGAGAUCAUCCGC-3’. Oligonucleotides for human Drp1siRNA were synthesized by IDT Oligo against target sequence 5’-GCCAGCUAGAUAUUAACAACAAGAA-3’. As a control, Silencer Negative Control 5’-CGUUAAUCGCGUAUAAUACGCGUAT-3’ (Ambion) was used.

### Antibodies

Anti-Mff (ProteinTech, 17090-1-AP) was used at 1:1000 dilution for western (WB) and 1:500 dilution for immunofluorescence (IF). Anti-Fis1 (ProteinTech, 10956-1-AP) was used at 1:1000 for WB and 1:500 for IF. Anti-Tubulin (DM1-α, Sigma/Aldrich) was used at 1:10,000 dilution for WB. Drp1 was detected using a rabbit monoclonal antibody (D6C7, Cell Signaling Technologies) at 1:500 dilution for IF. Anti-INF2 rabbit polyclonal was described previously (Ramabhadran et al., 2011). Organelle marker antibodies for WB include: anti-ATP synthase mouse monoclonal (Molecular Probes A21351), anti-Sec63 (Aviva ARP46839), and anti-Pmp70 rabbit polyclonal (Sigma 4200181), all used at 1:1000.

### Cell culture, transfection

Human osteosarcoma U2OS cells (American Type Culture Collection HTB96) were grown in DMEM (Invitrogen) supplemented with 10% calf serum (Atlanta Biologicals). Human PEX3-deficient fibroblasts (PBD400-T1) were a kind gift from Heidi McBride (Montreal Neurological Institute) and were grown in DMEM supplemented with 10% fetal calf serum (Atlanta Biologicals) and non-essential amino acids (GIBCO). To make the GFP-Drp1 KI U2OS cell line by CRISPR-Cas9, we used the GeCKO system (Zhang laboratory, MIT, http://genome-engineering.org/gecko/). The donor plasmid contained eGFP (A206K mutant) flanked by 445 bases upstream of hDrp1 start codon and 308 bases downstream from start (synthesized by IDT). The target guide sequence (CATTCATTGCCGTGGCCGGC) was predicted using the GeCKO website program and made by IDT. Donor and guide plasmids were transfected into U2OS cells at a 3:1 molar ratio using Lipofectamine 2000 (Invitrogen). Cells were put under puromycin selection and clones were selected by FACS sorting and single cell cloning, then verified by IF and Western blotting.

For transfection of the U2OS or Drp1 KI lines, cells were seeded at 4x10^5^ cells per well in a 6-well dish ~16 hours prior to transfection. Plasmid transfections were performed in OPTI-MEM media (Invitrogen) with 2 μL Lipofectamine 2000 (Invitrogen) per well for 6 hours, followed by trypsinization and re-plating onto concanavalin A (ConA, Sigma/Aldrich, Cat. No. C5275)-coated glass bottom MatTek dishes (P35G-1.5-14-C) at ~3.5x10^5^ cells per well. Cells were imaged in live cell media (Life Technologies, Cat.No. 21063-029), ~16-24 hours after transfection.

For all experiments, the following amounts of DNA were transfected per well (individually or combined for co-transfection): 500 ng for mito-BFP, eBFP2-Peroxisome, and mCherry-mito7; 850 ng for Tom20-mCherry; 1000 ng for ER-tagRFP, mPlum-mito3, and pEF.myc.ER-E2-Crimson; 100 ng for GFP-Mff-S, mStrawberry-Mff-S, GFP-Mff-S, and GFP-Fis1; 50 ng for MiD51-mStrawberry; 30 ng for mStrawberry-Rab4b and mStrawberry-Rab7a. 1000ng for AKAP1-FKBP12, and 500ng for Sac1-FKBP12 and GFP-Mff-FRB.

For siRNA transfections, cells were plated on 6 well plates with 30-40% density, and 2 μl RNAimax (Invitrogen) and 63 pg of siRNA were used per well. Cells were analyzed 72-84 hour post-transfection for suppression.

### Live imaging by confocal and Airyscan microscopy

Cells were grown on glass bottom matTek dishes coated with ConA (coverslips treated for ~2 hours with 100 μg/mL ConA in water at room temperature). MatTek dishes were loaded to a Wave FX spinning disk confocal microscope (Quorum Technologies, Inc., Guelph, Canada, on a Nikon Eclipse Ti microscope), equipped with Hamamatsu ImageM EM CCD cameras and Bionomic Controller (20/20 Technology, Inc) temperature-controlled stage set to 37°C. After equilibrating to temperature for 10 min, cells were imaged with the 60x 1.4 NA Plan Apo objective (Nikon) using the 403 nm laser and 450/50 filter for BFP, 491 nm and 525/20 for GFP, 561 nm and 593/40 for mStrawberry or mCherry, and 640 nm and 700/60 for mPlum and E2-Crimson. For rapamycin induction, cells were treated with freshly prepared rapamycin (Fisher Scientific, 10 mM Stock in DMSO, 10 μM final concentration on cells) during imaging.

Airyscan images were acquired on LSM 880 equipped with 63x/1.4 NA plan Apochromat oil objective, using the Airyscan detectors (Carl Zeiss Microscopy, Thornwood, NY). The Airyscan uses a 32-channel array of GaAsP detectors configured as 0.2 Airy Units per channel to collect the data that is subsequently processed using the Zen2 software. After equilibrating to 37 °C for 30 min, cells were imaged with the 405 nm laser and 450/30 filter for BFP, 488 nm and 525/30 for GFP, 561 nm and 595/25 for mStrawberry or mCherry, and 633 nm and LP 625 for mPlum.

### Immunofluorescence staining

Cells were fixed with 4% formaldehyde (Electron Microscopy Sciences, PA) in phosphate-buffered saline (PBS) for 10 min at room temperature. After washing with PBS three times, cells were permeabilized with 0.1% Triton X-100 in PBS for 15 min on ice. Cells were then washed three times with PBS, blocked with 0.5% BSA in PBS for 1 hour, and incubated with primary antibodies in diluted blocking buffer overnight. Wash with PBS three times. Mff or Fis1 polyclonal antibodies (Rabbit) were conjugated to Alexa Fluor 488 while PMP70 antibodies (Rabbit) were conjugated to Alexa Fluor 647 (Zenon Tricolor Rabbit IgG1 Labeling Kit, Molecular Probes, Invitrogen); Secondary antibodies were applied for 1hr at room temperature. After washing with PBS three times, samples were mount on vectashield (Vector lab. H-1000).

## Image analysis

### ER association of Drp1, Mff and Fis1

Cells expressing GFP-Drp1, Mff or Fis1, and markers for ER, mitochondria and peroxisomes, were imaged in a single focal plane for three min at 1.5-2 sec intervals. Regions of cells where tubular ER could be readily resolved and appeared continuous in a single plane of view were analyzed. Independent Drp1, MFF, or Fis1 punctae were counted as always associated if they remained in contact with the ER during every frame of the video, sometimes ER associated if the independent punctae contacted the ER at least half of total frames where punctae are visible, and not ER associated if no ER contact was visible.

### Drp1 punctae quantification

Drp1 KI cells transiently transfected with mitochondrial markers were imaged live by spinning disc confocal fluorescence microscopy for 10 min at 3 sec intervals in a single focal plane. Regions of interest with readily resolvable mitochondria and Drp1 were processed as described previously (Ji et al., 2015). We thresholded mitochondrially associated Drp1 punctae by using the Colocalization ImageJ plugin with the following parameters: Ratio 50%(0-100%); Threshold channel 1: 30 (0-255); Threshold channel 2: 30 (0-255); Display value: 255 (0-255). Mitochondrially associated Drp1 punctae were further analyzed by Trackmate as described previously (Ji et al., 2015). The number of Drp1 punctae were automatically counted frame-by-frame using the Find Stack Maxima ImageJ macro. The density of independent Drp1 punctae was quantified by visual assessment of each Drp1 puncta in an ROI for association with the mitochondria or peroxisome marker. Those punctae associated with neither mitochondria or peroxisomes were classified as independent. The result is expressed as number of independent Drp1 punctae per area of the ROI in square microns.

### Mitochondrial division rate

Described in detail in (Ji et al., 2015). Suitable ROI’s were selected for analysis based on whether individual mitochondria were resolvable and did not leave the focal plane. Files of these ROIs were assembled, then coded and scrambled by one investigator, and analyzed for division by a second investigator in a blinded manner as to the treatment condition. The second investigator scanned the ROIs frame-by-frame manually for division events, and determined total mitochondrial length within the ROI using the ImageJ macro, Mitochondrial Morphology. The results were then given back to the first investigator for de-coding. Division rate was analyzed over a 10-min period after DMSO, ionomycin (4 μM) or rapamycin (10 μM) treadment, depending on the experiment.

### Cell Fractionation

Modification of method in (Clayton and Shadel, 2014). All protease inhibitors from EMD Chemicals. For U20S, cells (12 x 75 cm^2^ flasks grown to approximately 70% confluence) were harvested by trypsinization and washed 3x with PBS. Post-trypsinization, all steps conducted at 4°C or on ice. The cell pellet (approximately 0.2 mL) was resuspended in 5.4 mL hypotonic buffer (10 mM Tris-HCl pH 7.5, 10 mM NaCl, 1.5 mM MgCh, protease inhibitors (2 μg/mL leupeptin, 10 μg/mL aprotinin, 2 μg/mL pepstatin A, 5 μg/mL calpain inhibitor 1, 5 μg/mL calpeptin, 1 mM benzamidine, 0.05 μg/mL cathepsin B inhibitor II), incubated for 10 min and lysed by dounce (Wheaton Dura-Grind), followed by addition of 3.6 mL 2.5x isotonic buffer (525 mM mannitol, 175 mM sucrose, 12.5 mM Tris-HCl pH 7.5, 2.5 mM EDTA, protease inhibitors). The lysate was centrifuged at 1300xg for 5 min (low-speed centrifugation). The low-speed supernatant was centrifuged at 13,000xg for 15 min (medium-speed centrifugation). The medium-speed supernatant was centrifuged at 208,000xg for 1 hr (high-speed centrifugation). For PEX3-deficient fibroblasts, conditions were similar except that four centrifugation speeds were used, following (Sugiura et al., 2017): 800xg for 10 min (nuclei and un-lysed cells, discarded), 2300xg (low-speed centrifugation), 23,000xg (medium-speed centrifugation), and 208,000xg for 1 hr (high-speed centrifugation). All pellets were washed with 1x isotonic buffer then resuspended in SDS-PAGE buffer. For sucrose gradient fractionation, the medium-speed supernatant (3.4 mL) was layered onto a discontinuous gradient containing equal volumes (1.9 mL) of 0.5, 0.75, 1, and 1.3 M sucrose (all in the background of 1x isotonic buffer), and centrifuged for 1 hr at 35,000 rpm in an SW41 rotor (Beckman Coulter) with no brake. Fractions (1 mL) were removed from top.

### Western blotting

Cells were grown on 6 well plate, trypsinized, washed with PBS and resuspended 50 μL PBS. This solution was mixed with 34 μL of 10% SDS and 1 μL of 1 M DTT, boiled 5 minutes, cooled to 23°C, then 17 μ1 of 300 mM of freshly made NEM in water was added. Just before SDS-PAGE, the protein sample was mixed 1:1 with 2xDB (250 mM Tris-HCl pH 6.8, 2 mM EDTA, 20% glycerol, 0.8% SDS, 0.02% bromophenol blue, 1000 mM NaCl, 4 M urea). Proteins were separated by 7.5% SDS-PAGE and transferred to a polyvinylidine difluoride membrane (Millipore). The membrane was blocked with TBS-T (20 mM Tris-HCl, pH 7.6, 136 mM NaCl, and 0.1% Tween-20) containing 3% BSA (Research Organics) for 1 hour, then incubated with the primary antibody solution at 4°C overnight. After washing with TBS-T, the membrane was incubated with horseradish peroxidase (HRP)-conjugated secondary antibody (Bio-Rad) for 1 hour at room temperature. Signals were detected by Chemiluminescence (Pierce). For western blotting of Mff KO cells, the Li-Cor Odyssey CLx system was used (Li-Cor Biotechnology, Lincoln NE), as well as IRDye-labeled anti-Rabbit and anti-mouse secondary antibodies from the same company.

## Acknowledgements

We would like to thank Mike Ryan for GFP-Fis1, Ayumu Sugiura for advice on organelle fractionation, Heidi McBride for PEX3-deficient cells, Maya Schuldiner for discussions on TA proteins, Anna Hatch for editing, and Peter Rocs for being receptive to anything. Supported by NIH GM069818 and GM106000 to HNH, NIH NS056244 and NS 087908 to SS, NIH P20 GM113132 to the BioMT COBRE and NIH S10OD010330 to the Norris Cotton Cancer Center.

The authors declare no competing financial interests.

---

**Figure S1.**
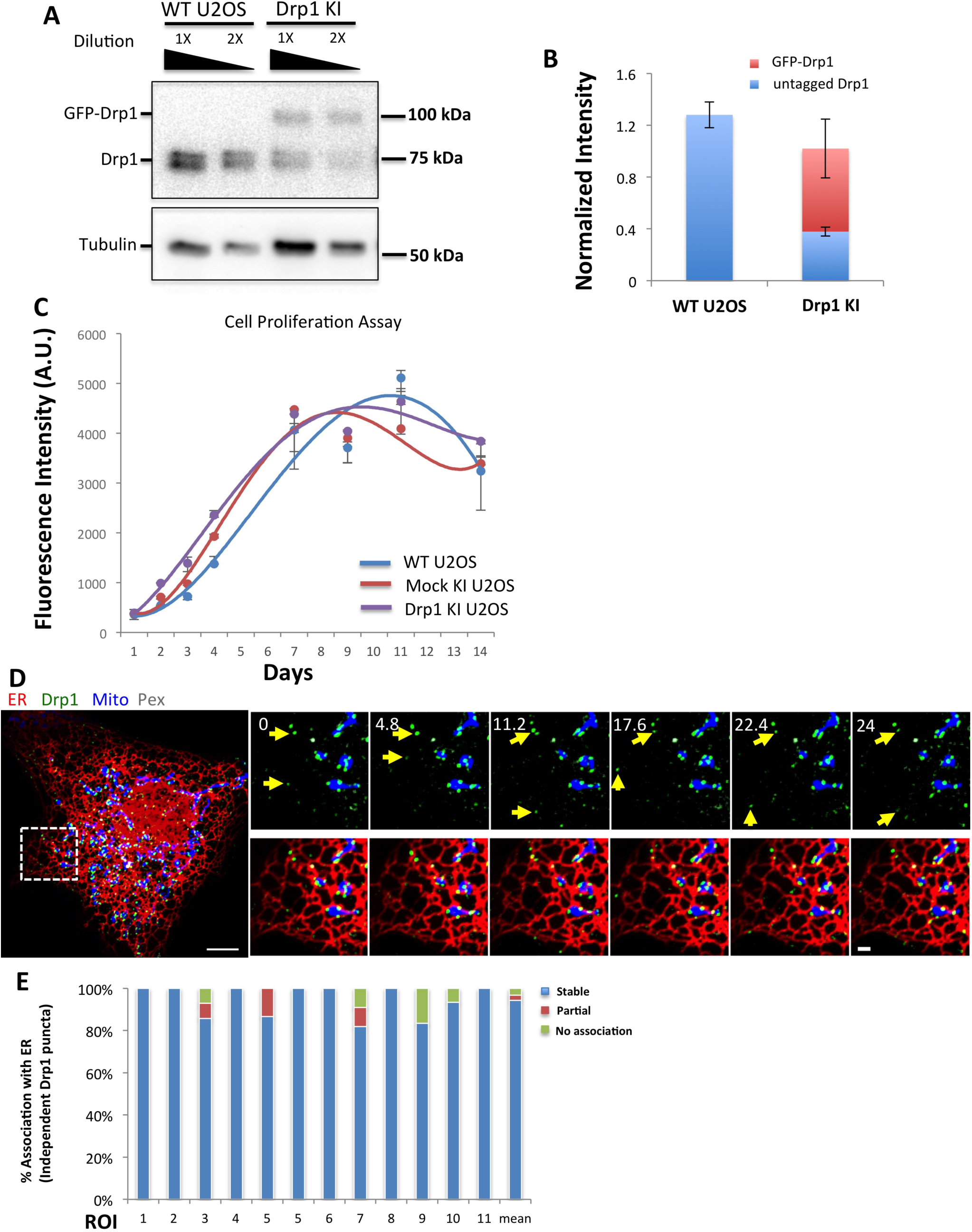
Characterization of Drp1 KI U2OS cell, and Drp1-independent punctae in Cos7 cells. (A) Western blot of U2OS cells and Drp1 KI U2OS cells showing expression level of GFP-Drp1 and untagged Drp1 with two dilutions of extract loaded (1x and 2x dilution). (B) Quantification of un-tagged Drp1 and GFP-Drp1 in WT U2OS and Drp1 KI cells from western blots (normalized to tubulin level). Error bars, S.D. (C) Cell proliferation assay (Alamar blue). Three replicates taken for each time point (median shown, with error bars representing minimum and maximum). Starting density: 5000 cells/24-well plate. Representative result from two independent experiments. (D) Drp1-independent punctae in Cos7 cells. *Left:* merged image of a live COS7 cell transiently expressing mito-BFP (mitochondria, blue), eBFP2-PMP20 (peroxisome, gray), GFP-Drp1(green) and ER-TagRFP (ER, red). *Right:* insets from boxed region. Yellow arrows denote independent Drp1 punctae associating with ER. (E) Graph depicting the degree of association between independent Drp1 punctae and ER in Cos7 cells, during 3 minute movies imaged every 1.5 sec. 175 independent Drp1 puncta were analyzed from 12 ROIs from 12 cells as shown in A. Stale association 94.2%±7.6%; Partial association 2.5%±4.7%; No association 3.3%±5.4%. Scale bar, 10 μm in whole cell image; 2 μm in inset. Time in sec.

**Figure S2.**
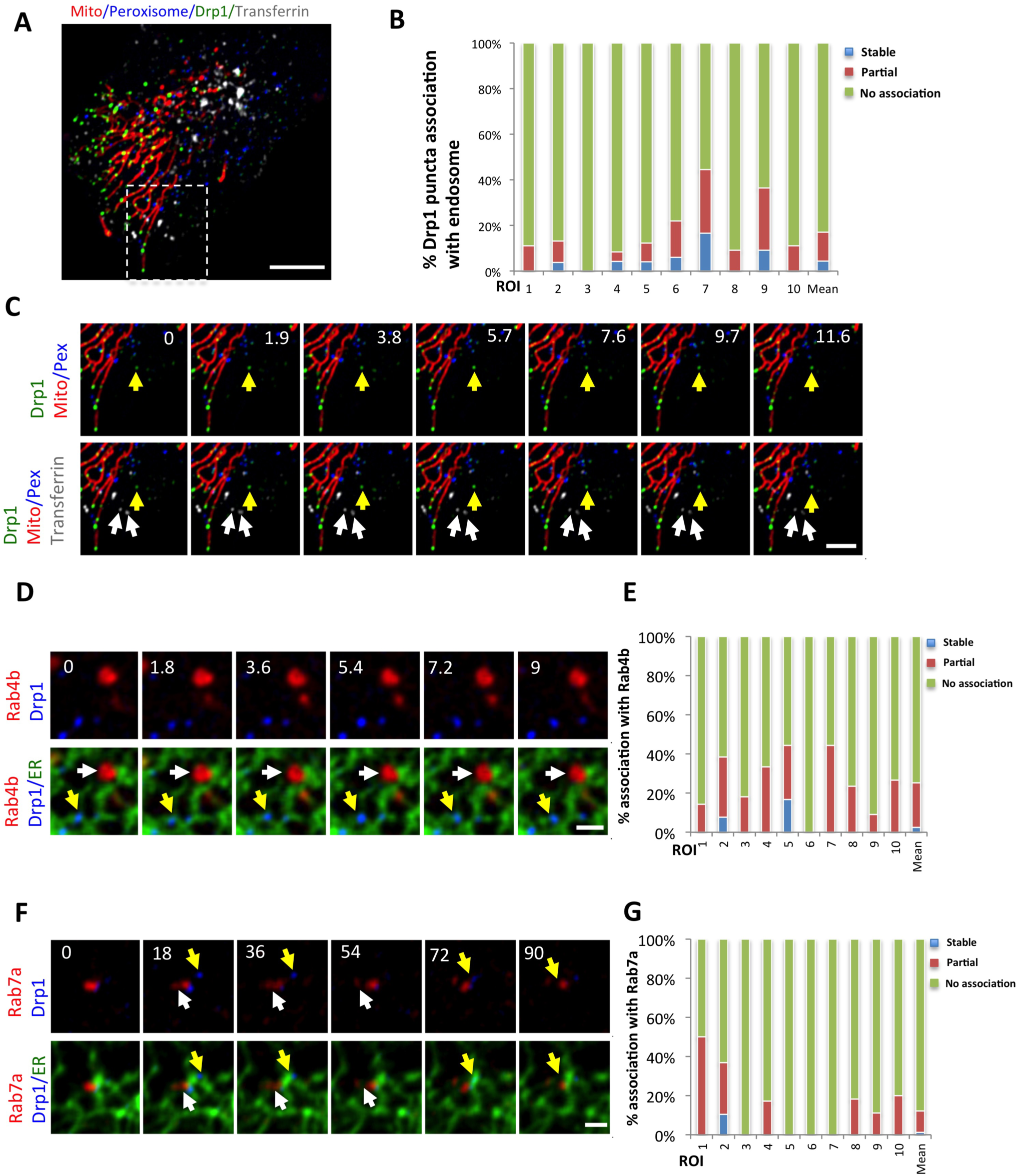
Independent Drp1 punctae are not stably associated with endosomal membranes. (A) Merged image of GFP-Drp1-KI cell expressing mCherry-mito3 (red); eBFP2-peroxisome (blue); and treated with transferrin-Alex647 (endosomes, gray). GFP-Drp1 in green. (B) Graph depicting degree of association between independent Drp1 punctae and transferrin-labeled membranes during 3 min videos imaged every 1.7 sec. 10 ROIs from 10 cells, 342 Drp1 punctae. (C) Time-lapse of inset from (A) showing independent Drp1 punctae distinct from transferrin-labeled endosomes. White arrows denote endosomes, yellow arrow denotes independent Drp1 puncta. (D) Time-lapse ROI of a GFP-Drp1-KI cell expressing mStrawberry-Rab4b (red), and ER-eBFP2 (ER, green). Drp1 in blue. Arrows defined as in C. (E) Graph depicting the degree of association between independent Drp1 punctae and Rab4b-labeled membranes during 3 min videos imaged every 1.8 sec. 10 ROIs from 10 cells, 152 Drp1 punctae. (F) Time-lapse of a Drp1 KI cell expressing mStrawberry-Rab7a (red),and ER-eBFP2 (ER, green). GFP-Drp1 is blue. Arrows defined as in C. (G) Graph depicting the degree of association between independent Drp1 punctae and Rab7a labeled membranes during 3 min videos imaged every 1.5 sec. 10 ROIs from 10 cells, 177 Drp1 punctae. Scale bar, 10 μm in (A); 5 μm in (C); 2 μm in (D) and (F). Time in sec.

**Figure S3.**
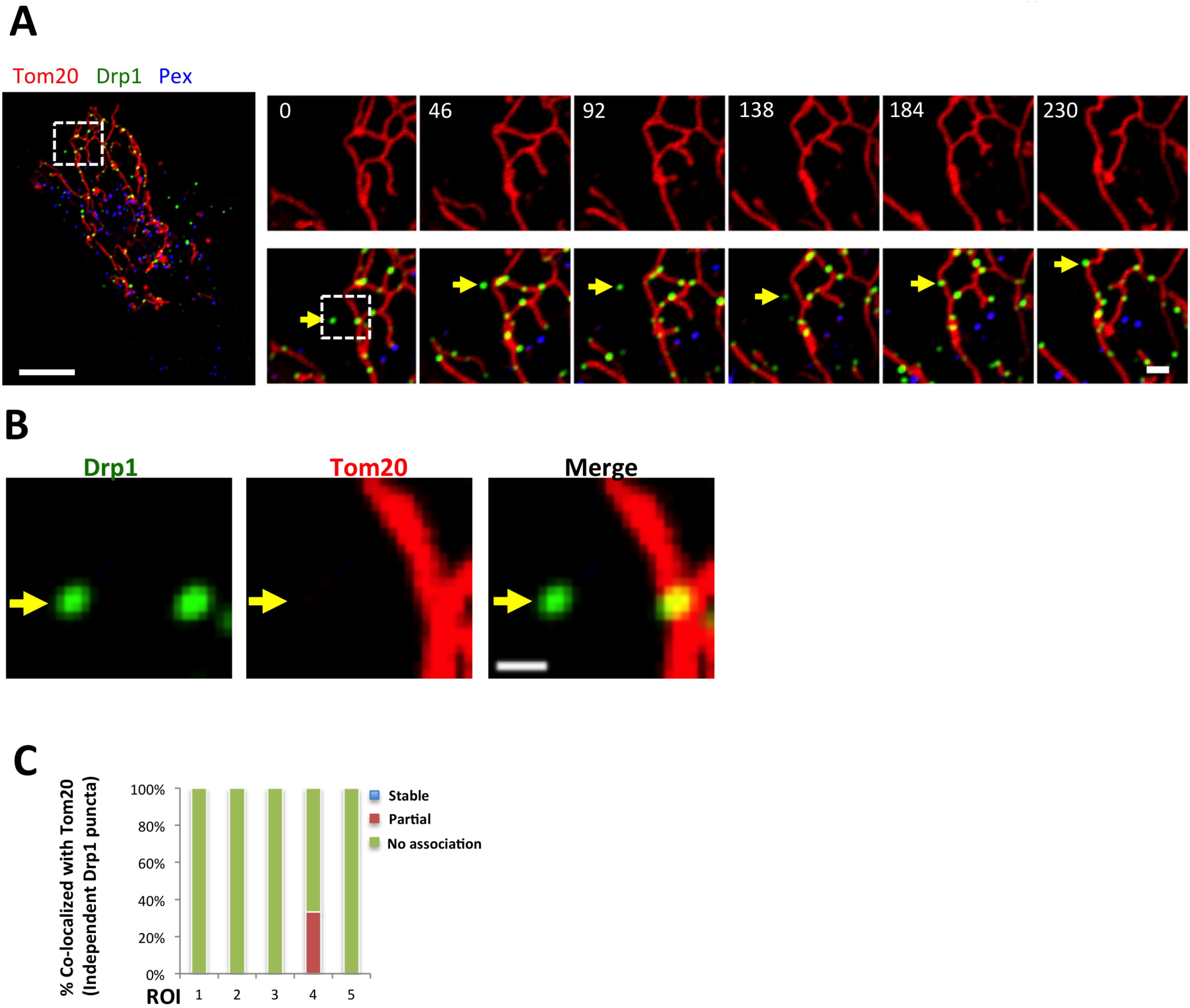
Independent Drp1 punctae do not associate with Tom20. (A) GFP-Drp1-KI cells transiently transfected with Tom20-mCherry (red) and eBFP2-PMP20 (peroxisomes, blue). Drp1 in green. Whole cell overlay on left, and time course of the indicated ROI on right (top, Tom20 alone. Bottom, merged image). (B) Zoom of indicated region of 0 sec time point in A, showing Drp1 and Tom20. No peroxisomes detected in this region. Yellow arrow denotes independent Drp1 puncta with no associated Tom20 signal. (C) Graph of percentage of independent Drp1 puncta overlaying with Tom20 (16 independent puncta from five ROI analyzed). One instance of overlap observed in ROI 4. Scale bar, 10 μm in whole cell in (A), 2 μm in inset in (A), 1 μm in (B). Time in sec.

**Figure S4.**
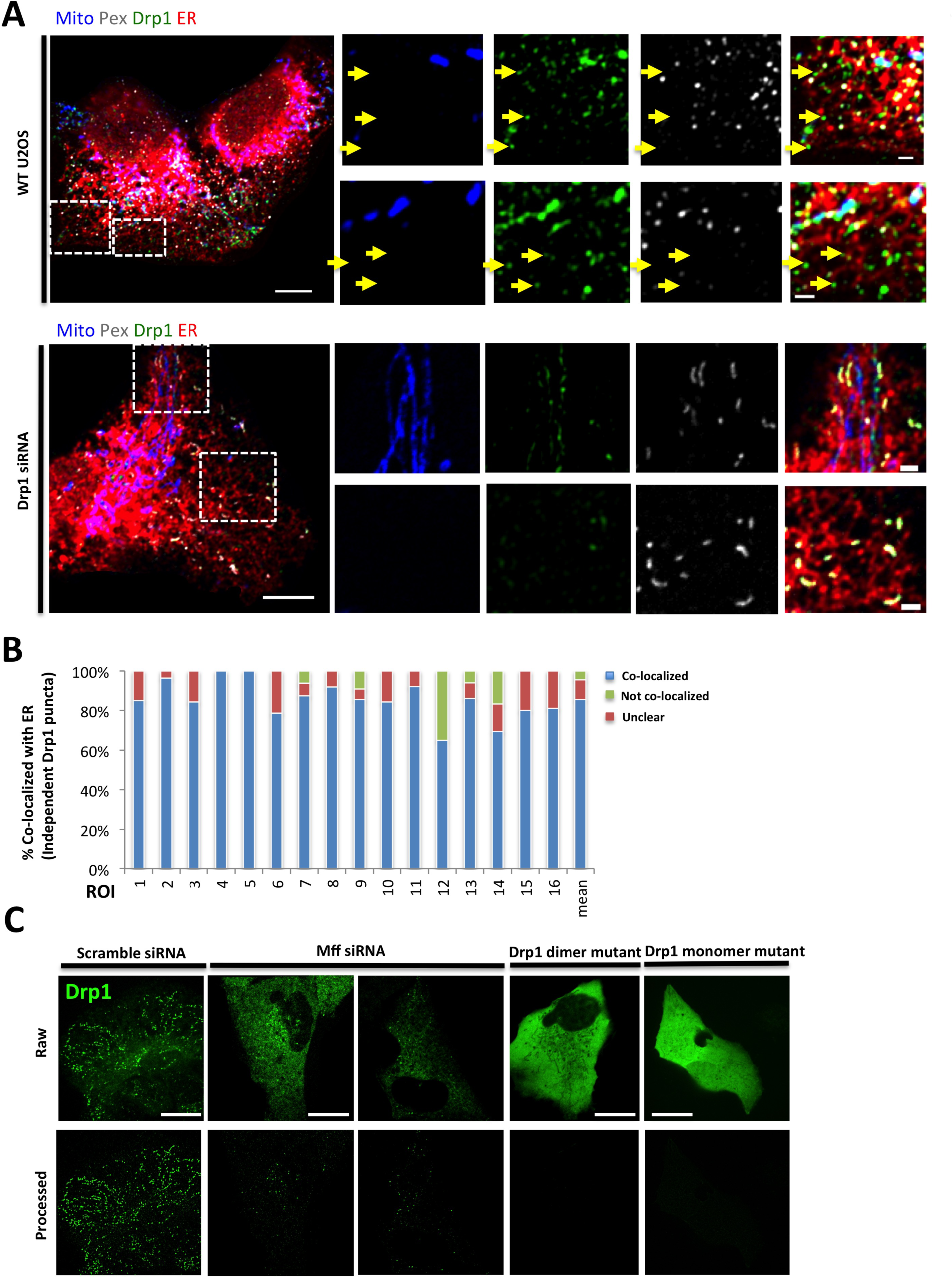
Experiments to test Drp1 aggregation. (A) Endogenous Drp1 staining by immunofluorescence in WT (top) and Drp1 KD U2OS cells (bottom). Also stained are mitochondria (blue), peroxisomes (gray) and ER (red). Yellow arrows indicate independent Drp1 puncta. WT and KD images acquired and processed identically. (B) Quantification of co-localization between endogenous Drp1 punctae and ER in WT U2OS cells. 562 independent puncta counted from 16 cells. Co-localized: 85.5%±9.7%; Not co-localized: 10.0%±7.3%; Unclear: 4.6%±9.4%. (C) Comparison of GFP-Drp1 distribution in U2OS cells under three conditions: GFP-Drp1-KI cells transfected with a scrambled siRNA (left), siRNA for Mff (center), and U2OS cells over-expressing GFP-Drp1 dimer or monomer mutant (right). Top row represents raw images and bottom row shows processed images to reveal Drp1 punctae, as described in methods (background subtracted and smoothed using imageJ). Scale bar, 10 μm in whole cell images in A and in C, 2μm in zoomed images in A.

**Figure S5.**
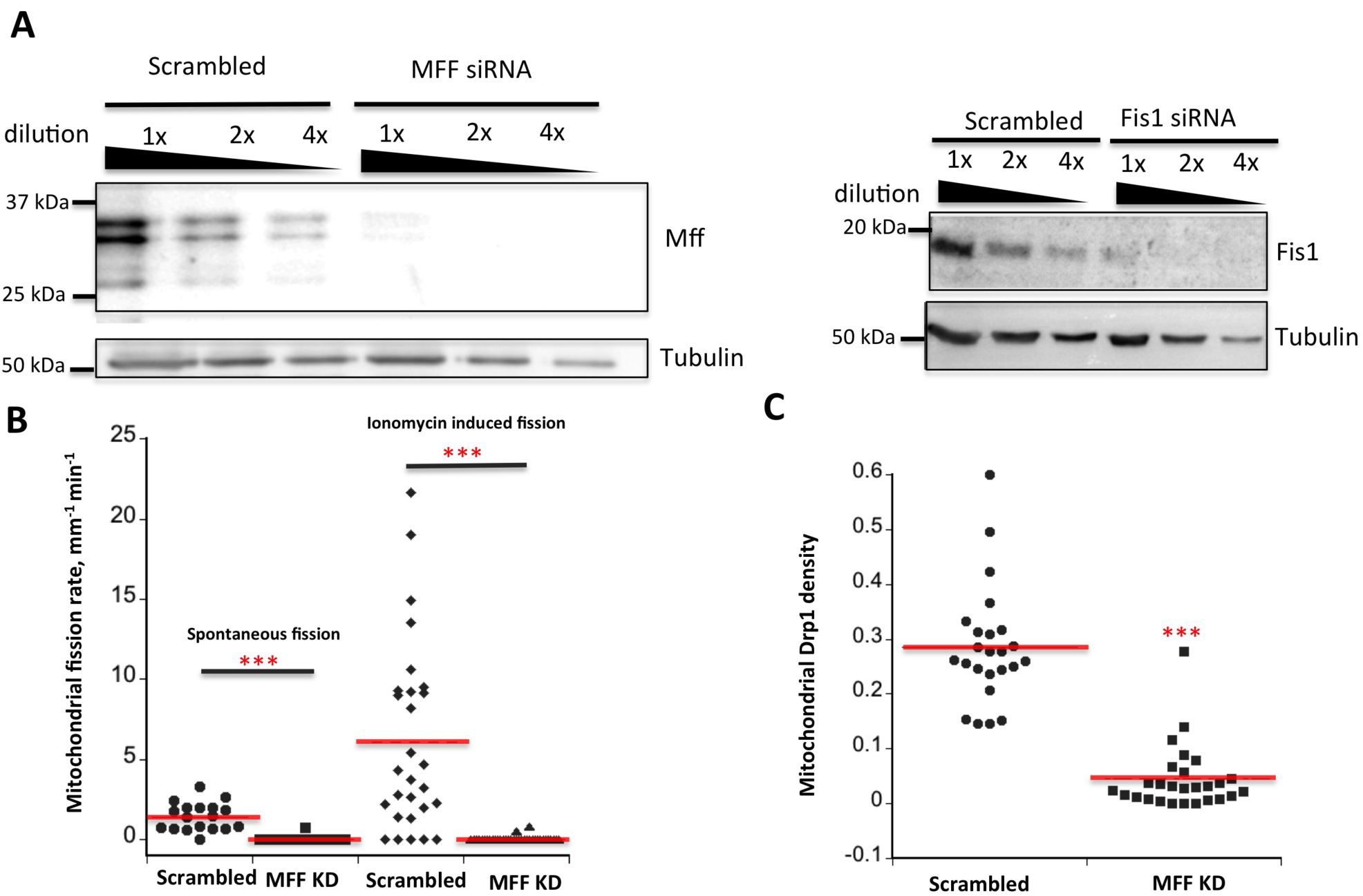
siRNA treatments for Mff and Fis1 in U2OS cells. (A) Western blots showing effectiveness of siRNA against Mff and Fis1. (B) Division rate quantification for scrambled siRNA and Mff siRNA in GFP-Drp1-KI U2OS cells, in both the unstimulated and ionomycin-stimulated states. In quantification of spontaneous division, 18 scrambled siRNA cells and 17 Mff siRNA cells are analyzed. In quantification of ionomycin induced division, 30 scrambled siRNA cells and 32 Mff siRNA cells are analyzed. ***, P<0.005, unpaired student t test. (C) Mitochondrial Drp1 punctae density quantification (units, Drp1 puncta per μm) for scrambled siRNA and Mff siRNA in GFP-Drp1-KI U2OS cells. 24 ROIs from 20 control cells and 27 ROIs from 25 MFF KD cells are analyzed. ***, P<0.005, unpaired student t test.

**Figure S6.**
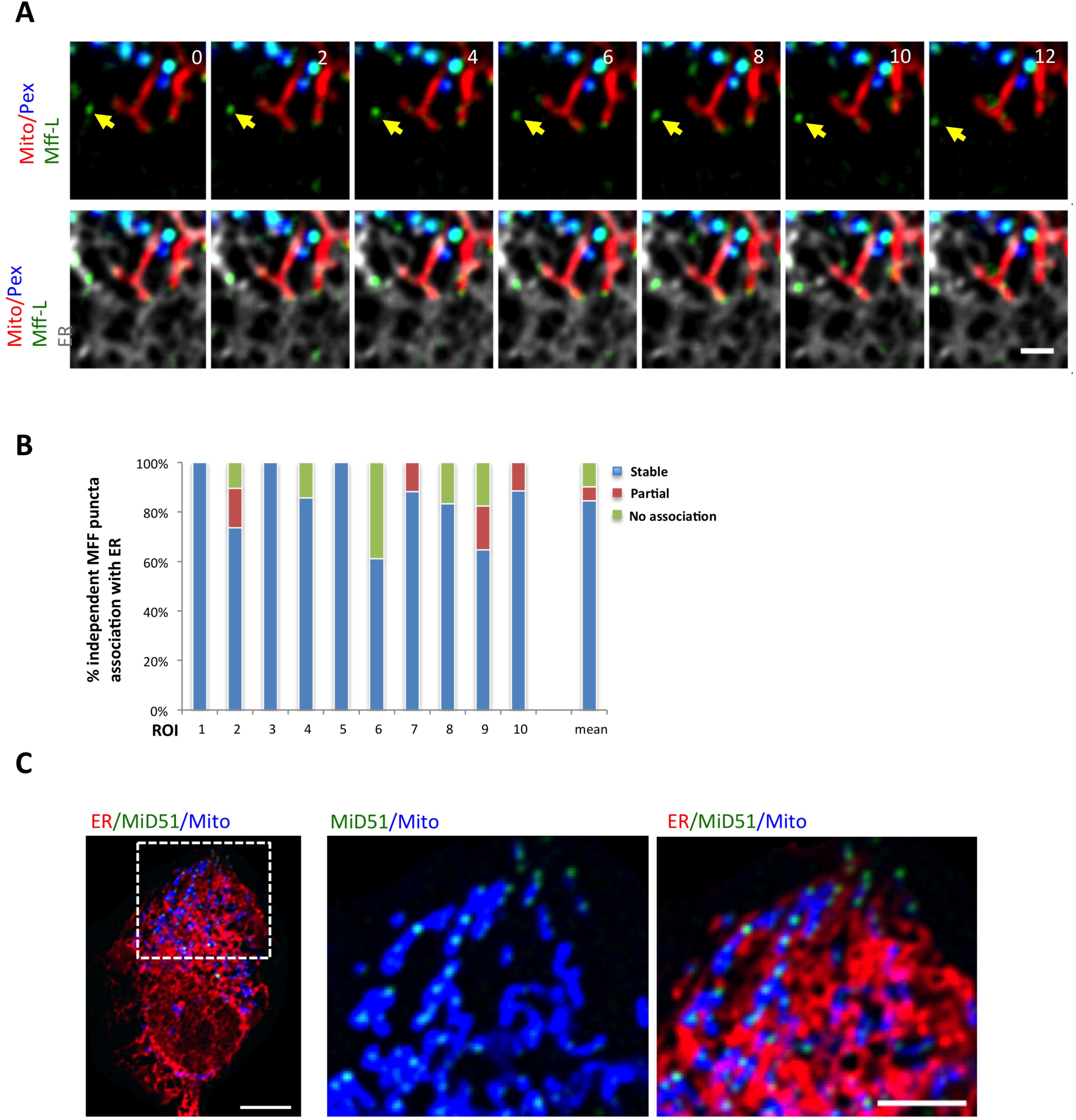
The Mff-L isoform displays ER-associated punctae, while Mid51 does not localize to ER. (A) Time-lapse from region of U2OS cell expressing mCherry-mito3 (red); eBFP2-peroxisome (blue); GFP-Mff-L (green); and E2-Crimson-ER (gray). Yellow arrow denotes independent Mff puncta associating with ER tubules. (B) Graph depicting the degree of association between independent Mff-L punctae and ER during 3 min videos imaged every 2 sec. 10 ROIs from 8 U2OS cells, 167 independent Mff punctae. (C) GFP-MiD51 does not display ER-associated punctae independent of mitochondria. Left panel: merged image of a live cell expressing MiD51-GFP (green), mitoBFP (blue) and ER-tagRFP (ER, red). *Right:* insets. Scale bar, 2 μm in A; 10 μm in whole cell in C, and 5 μm in inset of C. Time in sec.

## Video legends

**Video 1:** Confocal time-lapse of independent Drp1 puncta stably associating with ER tubules (yellow arrow) in GFP-Drp1-KI cell transiently expressing mPlum-mito3 (gray), eBFP2-PMP20 (blue) and ER-tagRFP (red). Drp1 in green. Left: without ER. Right: with ER. Time lapse taken in single z-plane every 1.77 sec. Time min:sec. Bar,2μm. (Fig. 1C)

**Video 2.** Confocal time-lapse of independent Drp1 puncta transferring to mitochondria in GFP-Drp1-KI U2OS cells transiently expressing mCherry-mito-7 (mitochondria, red), and eBFP2-PMP20 (peroxisome, blue). Time lapse taken in single z-plane in dorsal region of cell every 2.1 sec. Time min:sec. Bar, 2μm. (Fig. 2A)

**Video 3.** Confocal time-lapse of independent Drp1 puncta transferring from ER to mitochondria in GFP-Drp1-KI U2OS cells transiently expressing mito-BFP (Mito in red), mPlum-PMP20 (Pex in gray), and ER-tagRFP (ER in blue). Time lapse taken in single z-plane in dorsal region of cell every 3 sec. Left: Drp1 only. Middle: without ER. Right: with ER. Time min:sec. Bar, 2μm. (Fig. 2B)

**Video 4.** Confocal time-lapse of independent Drp1 puncta transferring to mitochondria, followed by mitochondrial division, in GFP-Drp1-KI U2OS cells transiently expressing mCherry-mito-7 (Mito in red), and eBFP2-PMP20 (Pex in blue). Time lapse taken in single z-plane in dorsal region of cell every 1.5 sec. Cells were treated with ionomycin (4 μM) to stimulate division at time 0. Time min:sec. Bar, 2μm. (Fig. 2C)

**Video 5.** Confocal time-lapse of independent Drp1 puncta transferring from ER to mitochondria, followed by mitochondrial division, in GFP-Drp1-KI U2OS cells transiently expressing mito-BFP (Mito in red), mPlum-PMP20 (Pex in gray), and ER-tagRFP (ER in blue). Time lapse taken in single z-plane in dorsal region of cell every 3 sec. Left: without ER. Right: with ER. Cells were treated with ionomycin (4 μM) to stimulate division at time 0. Time min:sec. Bar, 2μm. (Fig. 2D)

**Video 6:** Confocal time-lapse of independent Mff punctae on ER in U2OS cell transiently expressing mCherry-mito7 (gray), GFP-Mff-S (green), eBFP2-peroxisome (blue) and ER-E2-Crimson (Red). Time lapse was taken in single z-plane every 1.8 sec. Time min:sec. Bar, 2μm. (Fig. 4C)

**Video 7:** Airyscan time-lapse of independent Mff punctum transfer from ER to mitochondrion in U2OS cell transiently expressing mPlum-mito-3 (gray), GFP-Mff-S (green), eBFP2-PMP20 (blue) and ER-tagRFP (red). Left: without ER. Right: with ER. White arrow denotes independent Mff. Yellow arrow denotes peroxisome-associated Mff. Taken in single z-plane every 24 sec. Time min:sec. Bar, 2μm. (Fig. 6 A)

**Video 8:** Confocal time-lapse of Drp1 appearance and maturation at Mff-enriched site on ER in a GFP-Drp1-KI cell transiently expressing mito-BFP (blue), eBFP2-PMP20 (blue), mStrawberry-Mff-S (red), and ER-E2-Crimson (white). Drp1 in green. Left: Mff only. Middle: Drp1 only. Right: both Mff and Drp1. Time lapse was taken every 1.7 sec. Time min:sec. Bar, 2μm. (Fig. 7 A)

**Video 9:** Confocal time-lapse of Drp1 oligomerization upon ionomycin treatment in GFP-Drp1-KI U2OS cell transiently expressing mPlum-mito-3 (gray), eBFP2-PMP20 (blue) and ER-tagRFP (red). Drp1 in green. Left: Drp1 only. Right: Drp1 Mito Pex with ER. Taken in single z-plane in every 23 sec. Time min:sec. Ionomycin treatment at 1:30. Bar, 2μm. (Fig 9 A)

**Video 10**: Confocal time-lapse of Drp1 oligomerization after LatA (μM, 10min) pre-treatment followed by ionomycin treatment (4μM) in Drp1 KI U2OS cell transiently expressing mPlum-mito-3 (gray), eBFP2-peroxisome (blue) and ER-tagRFP (Red). Left: Drp1 only. Right: Drp1 Mito Pex with ER. Time lapse was taken in single z-plane in every 23.6 sec. Time min:sec. Ionomycin treatment at 1:34. Bar, 2μm. (Fig 9 B)

## References

Aviram, N., T. Ast, E.A. Costa, E.C. Arakel, S.G. Chuartzman, C.H. Jan, S. Hassdenteufel, J. Dudek, M. Jung, S. Schorr, R. Zimmermann, B. Schwappach, J.S. Weissman, and M. Schuldiner. 2016. The SND proteins constitute an alternative targeting route to the endoplasmic reticulum. Nature. 540:134–138.

Bui, H.T., and J.M. Shaw. 2013. Dynamin assembly strategies and adaptor proteins in mitochondrial fission. Current biology: CB. 23:R891–899.

Bustillo-Zabalbeitia, I., S. Montessuit, E. Raemy, G. Basanez, O. Terrones, and J. C. Martinou. 2014. Specific interaction with cardiolipin triggers functional activation of dynamin-related protein 1. PLOS One. 9(7): e102738.

Chang, C.R., and C. Blackstone. 2007. Drp1 phosphorylation and mitochondrial regulation. EMBO reports. 8:1088-1089; author reply 1089-1090.

Chang, C.R., and C. Blackstone. 2010. Dynamic regulation of mitochondrial fission through modification of the dynamin-related protein Drp1. Annals of the New York Academy of Sciences. 1201:34–39.

Chhabra, E.S., V. Ramabhadran, S.A. Gerber, and H.N. Higgs. 2009. INF2 is an endoplasmic reticulum-associated formin protein. Journal of cell science. 122:1430–1440.

Clayton, D.A., and G.S. Shadel. 2014. Isolation of mitochondria from tissue culture cells. Cold Spring Harbor protocols. 2014:pdb prot080002.

Cribbs, J.T., and S. Strack. 2007. Reversible phosphorylation of Drp1 by cyclic AMP-dependent protein kinase and calcineurin regulates mitochondrial fission and cell death. EMBO reports. 8:939–944.

Csordas, G., P. Varnai, T. Golenar, S. Roy, G. Purkins, T.G. Schneider, T. Balla, and G. Hajnoczky. 2010. Imaging interorganelle contacts and local calcium dynamics at the ER-mitochondrial interface. Mol Cell. 39:121–132.

De Vos, K.J., V.J. Allan, A.J. Grierson, and M.P. Sheetz. 2005. Mitochondrial function and actin regulate dynamin-related protein 1-dependent mitochondrial fission. Current biology: CB. 15:678–683.

Denic, V., V. Dotsch, and I. Sinning. 2013. Endoplasmic reticulum targeting and insertion of tail-anchored membrane proteins by the GET pathway. Cold Spring Harbor perspectives in biology. 5:a013334.

DuBoff, B., M. Feany, and J. Gotz. 2013. Why size matters - balancing mitochondrial dynamics in Alzheimer's disease. Trends in neurosciences. 36:325–335.

Duboff, B., J. Gotz, and M.B. Feany. 2012. Tau Promotes Neurodegeneration via DRP1 Mislocalization In Vivo. Neuron. 75:618–632.

Friedman, J.R., L.L. Lackner, M. West, J.R. DiBenedetto, J. Nunnari, and G.K. Voeltz. 2011. ER tubules mark sites of mitochondrial division. Science. 334:358–362.

Frohlich, C., S. Grabiger, D. Schwefel, K. Faelber, E. Rosenbaum, J. Mears, O. Rocks, and O. Daumke. 2013. Structural insights into oligomerization and mitochondrial remodelling of dynamin 1-like protein. The EMBO journal. 32:1280–1292.

Gandre-Babbe, S., and A.M. van der Bliek. 2008. The novel tail-anchored membrane protein Mff controls mitochondrial and peroxisomal fission in mammalian cells. Mol Biol Cell. 19:2402–2412.

Hatch, A.L., P.S. Gurel, and H.N. Higgs. 2014. Novel roles for actin in mitochondrial fission. Journal of cell science. 127:4549–4560.

Hatch, A.L., W.K. Ji, R.A. Merrill, S. Strack, and H.N. Higgs. 2016. Actin filaments as dynamic reservoirs for Drp1 recruitment. Mol Biol Cell. 27(20): 3109–3121.

Horie, C., H. Suzuki, M. Sakaguchi, and K. Mihara. 2002. Characterization of signal that directs C-tail-anchored proteins to mammalian mitochondrial outer membrane. Mol Biol Cell. 13(5): 1615–1625.

Hung, V., S.S. Lam, N.D. Udeshi, T. Svinkina, G. Guzman, V.K. Mootha, S.A. Carr, and A.Y. Ting. 2017. Proteomic mapping of cytosol-facing outer mitochondrial and ER membranes in living human cells by proximity biotinylation. eLife. 6.

Ji, W.K., A.L. Hatch, R.A. Merrill, S. Strack, and H.N. Higgs. 2015. Actin filaments target the oligomeric maturation of the dynamin GTPase Drp1 to mitochondrial fission sites. Elife. 4:e11553.

Kobayashi, S., A. Tanaka, and Y. Fujiki. 2007. Fis1, DLPl, and Pexllp coordinately regulate peroxisome morphogenesis. Experimental cell research. 313:1675–1686.

Koch, A., Y. Yoon, N.A. Bonekamp, M.A. McNiven, and M. Schrader. 2005. A role for Fis1 in both mitochondrial and peroxisomal fission in mammalian cells. Mol Biol Cell. 16:5077–5086.

Koch, J., and C. Brocard. 2012. PEXll proteins attract Mff and human Fis1 to coordinate peroxisomal fission. Journal of cell science. 125:3813–3826.

Korobova, F., T.J. Gauvin, and H.N. Higgs. 2014. A role for myosin II in mammalian mitochondrial fission. Current biology: CB. 24:409–414.

Korobova, F., V. Ramabhadran, and H.N. Higgs. 2013. An actin-dependent step in mitochondrial fission mediated by the ER-associated formin INF2. Science. 339:464–467.

Krumpe, K., I. Frumkin, Y. Herzig, N. Rimon, C. Ozbalci, B. Brugger, D. Rapaport, and M. Schuldiner. 2012. Ergosterol content specifies targeting of tail-anchored proteins to mitochondrial outer membranes. Mol Biol Cell. 23:3927–3935.

Labbe, K., A. Murley, and J. Nunnari. 2014. Determinants and functions of mitochondrial behavior. Annual review of cell and developmental biology. 30:357–39l.

Lam, S.K., N. Yoda, and R. Schekman. 2010. A vesicle carrier that mediates peroxisome protein traffic from the endoplasmic reticulum. Proceedings of the National Academy of Sciences of the United States of America. 107:21523–21528.

Lee, J.E., L.M. Westrate, H. Wu, C. Page, and G.K. Voeltz. 2016. Multiple dynamin family members collaborate to drive mitochondrial division. Nature. 540:139–143.

Lewis, S.C., L.F. Uchiyama, and J. Nunnari. 2016. ER-mitochondria contacts couple mtDNA synthesis with mitochondrial division in human cells. Science. 353:aaf5549.

Li, S., S. Xu, B.A. Roelofs, L. Boyman, W.J. Lederer, H. Sesaki, and M. Karbowski. 2015. Transient assembly of F-actin on the outer mitochondrial membrane contributes to mitochondrial fission. J Cell Biol. 208:109–123.

Liu, R., and D.C. Chan. 2015. The mitochondrial fission receptor Mff selectively recruits oligomerized Drp1. Mol Biol Cell. 26:4466–4477.

Loson, O.C., Z. Song, H. Chen, and D.C. Chan. 2013. Fis1, Mff, MiD49, and MiD51 mediate Drp1 recruitment in mitochondrial fission. Mol Biol Cell. 24:659–667.

Macdonald, P. J., N. Stepanyants, N. Mehrotra, J. A. Mears, X. Qi, H. Sesaki, and R. Ramachandran. 2014. A dimeric equilibrium intermediate nucleates Drp1 reassembly on mitochondrial membranes. Mol Biol Cell. 25(12): 1905–1915.

Manor, U., S. Bartholomew, G. Golani, E. Christenson, M. Kozlov, H. Higgs, J. Spudich, and J. Lippincott-Schwartz. 2015. A mitochondria-anchored isoform of the actin-nucleating spire protein regulates mitochondrial division. Elife. 4.

Mateja, A., M. Paduch, H.Y. Chang, A. Szydlowska, A.A. Kossiakoff, R.S. Hegde, and R.J. Keenan. 2015. Protein targeting. Structure of the Get3 targeting factor in complex with its membrane protein cargo. Science. 347:1152–1155.

Matsui, T., T. Itoh, and M. Fukuda. 2011. Small GTPase Rab12 regulates constitutive degradation of transferrin receptor. Traffic. 12:1432–1443.

Mishra, P., and D.C. Chan. 2016. Metabolic regulation of mitochondrial dynamics. J Cell Biol. 212:379–387.

Moore, A.S., Y.C. Wong, C.L. Simpson, and E.L. Holzbaur. 2016. Dynamic actin cycling through mitochondrial subpopulations locally regulates the fission-fusion balance within mitochondrial networks. Nature communications. 7:12886.

Nunnari, J., and A. Suomalainen. 2012. Mitochondria: in sickness and in health. Cell. 148:1145–1159.

Osellame, L.D., A.P. Singh, D.A. Stroud, C.S. Palmer, D. Stojanovski, R. Ramachandran, and M.T. Ryan. 2016. Cooperative and independent roles of the Drp1 adaptors Mff, MiD49 and MiD51 in mitochondrial fission. Journal of cell science. 129:2170–2181.

Otera, H., N. Miyata, O. Kuge, and K. Mihara. 2016. Drp1-dependent mitochondrial fission via MiD49/51 is essential for apoptotic cristae remodeling. J Cell Biol. 212:531–544.

Otera, H., C. Wang, M.M. Cleland, K. Setoguchi, S. Yokota, R.J. Youle, and K. Mihara. 2010. Mff is an essential factor for mitochondrial recruitment of Drp1 during mitochondrial fission in mammalian cells. J Cell Biol. 191:1141–1158.

Palmer, C.S., K.D. Elgass, R.G. Parton, L.D. Osellame, D. Stojanovski, and M.T. Ryan. 2013. Adaptor proteins MiD49 and MiD51 can act independently of Mff and Fis1 in Drp1 recruitment and are specific for mitochondrial fission. The Journal of biological chemistry. 288:27584–27593.

Pernas, L., and L. Scorrano. 2016. Mito-Morphosis: Mitochondrial Fusion, Fission, and Cristae Remodeling as Key Mediators of Cellular Function. Annual review of physiology. 78:505–531.

Pitts, K.R., Y. Yoon, E.W. Krueger, and M.A. McNiven. 1999. The dynamin-like protein DLP1 is essential for normal distribution and morphology of the endoplasmic reticulum and mitochondria in mammalian cells. Mol Biol Cell. 10:4403–4417.

Ramabhadran, V., F. Korobova, G.J. Rahme, and H.N. Higgs. 2011. Splice variant-specific cellular function of the formin NF2 in maintenance of Golgi architecture. Mol Biol Cell. 22:4822–4833.

Ramachandran, R. 2017. Mitochondrial dynamics: the dynamin superfamily and execution by collusion. Sem Cell Dev Biol. 17: 30365–30368.

Richter, V., A.P. Singh, M. Kvansakul, M.T. Ryan, and L.D. Osellame. 2015. Splitting up the powerhouse: structural insights into the mechanism of mitochondrial fission. Cellular and molecular life sciences: CMLS. 72:3695–3707.

Ruan, L., C. Zhou, E. Jin, A. Kucharavy, Y. Zhang, Z. Wen, L. Florens, and R. Li. 2017. Cytosolic proteostasis through importing of misfolded proteins into mitochondria. Nature. 543:443–446.

Schrader, M., J.L. Costello, L.F. Godinho, A.S. Azadi, and M. Islinger. 2016. Proliferation and fission of peroxisomes - An update. Biochimica et biophysica acta. 1863:971–983.

Schuldiner, M., S.R. Collins, N.J. Thompson, V. Denic, A. Bhamidipati, T. Punna, J. Ihmels, B. Andrews, C. Boone, J.F. Greenblatt, J.S. Weissman, and N.J. Krogan. 2005. Exploration of the function and organization of the yeast early secretory pathway through an epistatic miniarray profile. Cell. 123:507–519.

Schuldiner, M., J. Metz, V. Schmid, V. Denic, M. Rakwalska, H.D. Schmitt, B. Schwappach, and J.S. Weissman. 2008. The GET complex mediates insertion of tail-anchored proteins into the ER membrane. Cell. 134:634–645.

Shen, Q., K. Yamano, B.P. Head, S. Kawajiri, J.T. Cheung, C. Wang, J.H. Cho, N. Hattori, R.J. Youle, and A.M. van der Bliek. 2014. Mutations in Fis1 disrupt orderly disposal of defective mitochondria. Mol Biol Cell. 25:145–159.

Soubannier, V., P. Rippstein, B.A. Kaufman, E.A. Shoubridge, and H.M. McBride. 2012. Reconstitution of mitochondria derived vesicle formation demonstrates selective enrichment of oxidized cargo. PloS one. 7:e52830.

Stefanovic, S., and R.S. Hegde. 2007. Identification of a targeting factor for posttranslational membrane protein insertion into the ER. Cell. 128:1147–1159.

Stojanovski, D., O.S. Koutsopoulos, K. Okamoto, and M.T. Ryan. 2004. Levels of human Fis1 at the mitochondrial outer membrane regulate mitochondrial morphology. Journal of cell science. 117:1201–1210.

Sugiura, A., S. Mattie, J. Prudent, and H.M. McBride. 2017. Newly born peroxisomes are a hybrid of mitochondrial and ER-derived pre-peroxisomes. Nature. 542:251–254.

Toyama, E.Q., S. Herzig, J. Courchet, T.L. Lewis, Jr, O.C. Loson, K. Hellberg, N.P. Young, H. Chen, F. Polleux, D.C. Chan, and R.J. Shaw. 2016. Metabolism. AMP-activated protein kinase mediates mitochondrial fission in response to energy stress. Science. 351:275–281.

Vafai, S.B., and V.K. Mootha. 2012. Mitochondrial disorders as windows into an ancient organelle. Nature. 491:374–383.

van der Zand, A., I. Braakman, and H.F. Tabak. 2010. Peroxisomal membrane proteins insert into the endoplasmic reticulum. Mol Biol Cell. 21:2057–2065.

Yagita, Y., T. Hiromasa, and Y. Fujiki. 2013. Tail-anchored PEX26 targets peroxisomes via a PEX19-dependent and TRC40-independent class I pathway. J Cell Biol. 200:651–666.

Yoon, Y., E.W. Krueger, B.J. Oswald, and M.A. McNiven. 2003. The mitochondrial protein hFis1 regulates mitochondrial fission in mammalian cells through an interaction with the dynamin-like protein DLP1. Molecular and cellular biology. 23:5409–5420.

Yoon, Y., K.R. Pitts, S. Dahan, and M.A. McNiven. 1998. A novel dynamin-like protein associates with cytoplasmic vesicles and tubules of the endoplasmic reticulum in mammalian cells. J Cell Biol. 140:779–793.

Youle, R.J., and A.M. van der Bliek. 2012. Mitochondrial fission, fusion, and stress. Science. 337:1062–1065.

Zhou, C., B.D. Slaughter, J.R. Unruh, F. Guo, Z. Yu, K. Mickey, A. Narkar, R.T. Ross, M. McClain, and R. Li. 2014. Organelle-based aggregation and retention of damaged proteins in asymmetrically dividing cells. Cell. 159:530–542.

